# Histone modifications form a cell-type-specific chromosomal bar code that modulates and maintains patterns of gene expression through the cell cycle

**DOI:** 10.1101/2020.10.16.341446

**Authors:** John A. Halsall, Simon Andrews, Felix Krueger, Charlotte E. Rutledge, Gabriella Ficz, Wolf Reik, Bryan M. Turner

**Author notes:** Corresponding Authors: Professor Bryan Turner, Dr John Halsall, Chromatin and Gene Regulation Group, Institute of Cancer and Genomic Sciences, College of Medical and Dental Sciences, University of Birmingham, B15 2TT.

## Abstract

**Background:** Chromatin configuration influences gene expression in eukaryotes at multiple levels, from individual nucleosomes to chromatin domains several Mb long. Post-translational modifications (PTM) of core histones seem to be involved in chromatin structural transitions, but how remains unclear.

To explore this, we used ChIP-seq and two cell types, HeLa and lymphoblastoid (LCL) to define how changes in chromatin packaging through the cell cycle influence the distributions of three transcription-associated histone modifications, H3K9ac, H3K4me3 and H3K27me3.

**Results:** Chromosome regions (bands) of 10-50Mb, detectable by immunofluorescence microscopy of metaphase (M) chromosomes, are also present in G_1_ and G_2_. We show that they comprise 1-5Mb sub-bands that differ between HeLa and LCL but remain consistent through the cell cycle. The same sub-bands are defined by H3K9ac and H3K4me3, while H3K27me3 spreads more widely.

We found little change between cell cycle phases, whether compared by 5Kb rolling windows or when analysis was restricted to functional elements such as transcription start sites and topologically associating domains.

Only a small number of genes showed cell-cycle related changes: at genes encoding proteins involved in mitosis, H3K9 became highly acetylated in G_2_M, possibly because of ongoing transcription.

**Conclusions:** Modified histone isoforms H3K9ac, H3K4me3 and H3K27me3 exhibit a characteristic genomic distribution at resolutions of 1Mb and below that differs between HeLa and lymphoblastoid cells but remains remarkably consistent through the cell cycle. We suggest that this cell-type-specific chromosomal bar-code is part of a homeostatic mechanism by which cells retain their characteristic gene expression patterns, and hence their identity, through multiple mitoses.

## INTRODUCTION

Post-translational modifications of histone proteins have been closely linked to control of gene expression at multiple levels. Modifications, singly or in combination, often act by providing binding sites through which regulatory proteins can be targeted to specific regions of the genome, thus allowing regulation of genes or groups of genes [1]. This process can operate at a local level through a promoter or other relatively small region (several Kb) but can also bring about more widespread changes in packaging. For example, some modifications, such as H3 acetylated at lysine 9 (H3K9ac) and H3 trimethylated at lysine 4 (H3K4me3) both show sharply defined peaks at the promoters and TSS of active and potentially active genes [2], while the mono-methylated H3K4 is specifically enriched at enhancers [3]. Other modifications mark silent regions, including highly condensed facultative and centric heterochromatin (H3K27me3 and H4K20me3 respectively), and are generally much more widely spread, in some cases covering several Mb [4,5]. In most cases the structural and functional mechanisms that underpin these associations are still incompletely understood.

Recently, attention has been drawn to the ability of chromatin to undergo liquidliquid phase transitions (LLPS, [6]), a physical shift with the potential to bring about increased chromatin compaction over large genomic regions [7–10]. At the same time, transcriptionally active chromatin has been shown to form condensates via transcriptional coactivators and RNA pol II phosphorylation [11–13]. Intriguingly, at least in model systems, histone acetylation levels along with acetylation-binding bromodomain proteins, can be used to regulated LLPS [14]. The overall reduction in net change brought about by lysine acetylation is also likely to be influential.

Progression through the cell cycle provides a challenge for maintenance of patterns of transcription, particularly with respect to the dramatic changes in chromatin compaction that accompany passage through mitosis. Chromosome condensation leading up to metaphase disrupts the 3D organisation of the genome, with implications for the retention of cell-specific gene expression patterns. For example, interphase chromosomes are organised into topologically associating domains (TADs). These, large self-interacting chromosome regions bring together genes and genomic elements and are closely involved in their regulation [15,16]. Histone modifications mark or are confined by TADS, showing similar distribution within individual domains but significant differences between neighbouring domains [17]. But TADs are undetectable in mitotic chromosomes [18], raising the question of how their cell-type-specific regulatory properties are conserved. Mitotic chromatin condensation is also associated with, and may sometimes be responsible for, the expulsion of some transcription factors from mitotic chromatin [19]. Others, sometimes referred to as bookmarking factors, are selectively retained [20]. A recent comprehensive proteomic analysis has suggested that only a minority of transcription factors found on chromatin were depleted during mitosis and the regulatory landscape remained relatively unchanged [21].

Maintenance of histone PTM levels and distribution through the cell cycle is complex, both because different PTM are maintained as a dynamic steady-state balance through the actions of different modifying and demodifying enzymes, and because newly-assembled chromatin at S-phase must somehow re-establish regional modification patterns. Histone modifications carried on the parental histones are diluted into newly formed chromatin and it is thought they act as template to reinstate chromatin marking to prereplication levels [22]. This happens at different rates for different modifications. Most modifications, including acetylation, are reinstated rapidly by the modification of new histones within a single cell cycle. However, the propagation of heterochromatic marks H3K27me3 and H3K9me3 on new histones takes longer than a cell cycle, requiring ongoing modification of old and new histones to maintain modification levels [23].

Overall levels of some histone PTM change rapidly as cells prepare to enter, or leave, mitosis. For example, histone H3 phosphorylation at serine 10 (H3S10ph) is dramatically increased during mitosis and seems to be essential for the compaction of chromatin required for cell division [24–26]. Phosphorylation of serine 28 [27] and threonine 3 [28] is also enriched in mitosis. It is interesting that all three phosphorylatable serines are adjacent to lysines whose modifications play critical roles in gene expression [29].

Histone acetylation levels are reduced during mitosis [21,30], and this deacetylation seems to be required for accurate chromatin segregation [31]. Nevertheless, we and others have shown that metaphase chromosomes contain readily detectable levels of the acetylated isoforms of all four core histones [30,32] and Ginno et al. showed relatively little change in chromatin accessibility (by ATAC-seq) even in mitotic chromatin [21]. Further, PTM associated with regulation of transcription (such as H3K9ac, H3K4me3, H3K27me3) are depleted in centromeric regions [30] but have characteristic and overlapping distributions (bands) across chromosome arms [32]. These bands, range in size from 10-50Mb and correspond to more gene-rich regions of the genome [32–34]. These PTM serve as markers for such regions through mitosis.

Here we use cell cycle sorting and ChIP-seq to analyse the genomic distribution of three histone modifications closely associated with transcriptional regulation (H3K9ac, H3K4me3 and H3K27me3) through the cell cycle in two human cell types. Reconstituting these data at pseudo-microscopic resolution we demonstrate that the banding patterns visible in mitotic chromosomes are remarkably strongly conserved from G_1_ to G_2_M and correlate with the patterns observed by metaphase chromosome staining. At higher (<5Mb) resolution, the bands were made up of clearly defined sub-bands that differed between celltypes. Remarkably, these patterns remained consistent through the cell cycle. At the individual gene level, we have identified a small number of local, cell-cycle-dependent changes within this stable, cell-specific landscape. These mark genes with mitosis-specific functions, whose selective transcription persists into G_2_M.

## RESULTS

Human lymphoblastoid cells (LCL, diploid, non-cancerous) and HeLa cells (aneuploid, cancer-derived) were fractionated into cell cycle phases (G_1_, G_2_M and M) by either flow cytometry (based on DNA content), centrifugal elutriation (largely based on size) or mitotic shake off (applicable only to HeLa). Details of cell culture conditions, fractionation techniques and the purity of isolated fractions are presented in Methods and supplementary figure S1. Overall, based on DNA content and microscopy, G_1_ fractions were 80-90% pure, G_2_M fractions were 70-90% pure and mitotic shake off fractions were almost 100% pure. G_2_M fractions were >90% G_2_ (ie. <10% M).

Fractionated cells were fixed in acetone prior to preparation of chromatin by micrococcal nuclease digestion. This revised procedure rapidly inactivates enzymes that might disrupt histone modifications during in-vitro processing while allowing efficient chromatin digestion to give nucleosomal fragments. We found no significant differences in chromatin yield or fragment size between different cell cycle fractions (Supplementary Figure S1E).

Chromatin was immunoprecipitated with rabbit polyclonal antisera to two histone modifications associated with transcriptionally active chromatin, H3 trimethylated at lysine 4 (H3K4me3), H3 acetylated at lysine 9 (H3K9ac) and one modification associated with gene silencing and formation of facultative heterochromatin, H3 tri-methylated at lysine 27 (H3K27me3). Overall, ChIP-seq data obtained using our protocol for LCL cells correlate closely with results for the same cell type available through ENCODE (supplementary figure S2).

### Bands of histone PTM detected by immunofluorescence microscopy on metaphase chromosomes are also present in interphase and are made up of smaller sub-bands

To define the regional distribution of the chosen modifications, ChIP-seq data were quantified using rolling windows and visualised using a red-green colour scale, mimicking metaphase chromosome staining by immunofluorescence (IF) (see Methods for details). These interphase chromosomes assembled from ChIP-seq data were aligned with the corresponding chromosome from LCL metaphase chromosome spreads, immunolabelled with antisera to H3K9ac, H3K4me3 and H3K27me3.

Presented in this way, H3K9ac and H3K4me3 show very similar banded distributions along chromosomes, with interspersed regions of relatively high (green) and low (red) modification (Figure 1A, complete ChIP-seq karyotypes in supplementary Figures S3 and S4). In the images shown, the pixel resolution (ie. minimum visible band size), is approximately 1.5Mb. Large regions (10-50Mb) relatively rich in modified histone and corresponding to bands detected by IF [32] were identifiable in interphase chromosomes as clusters of smaller (1-5Mb), enriched (green) bands. (Chromosomes 1, 3, 6 and 12 provide particularly clear examples, Figure 1).

**Figure 1:**
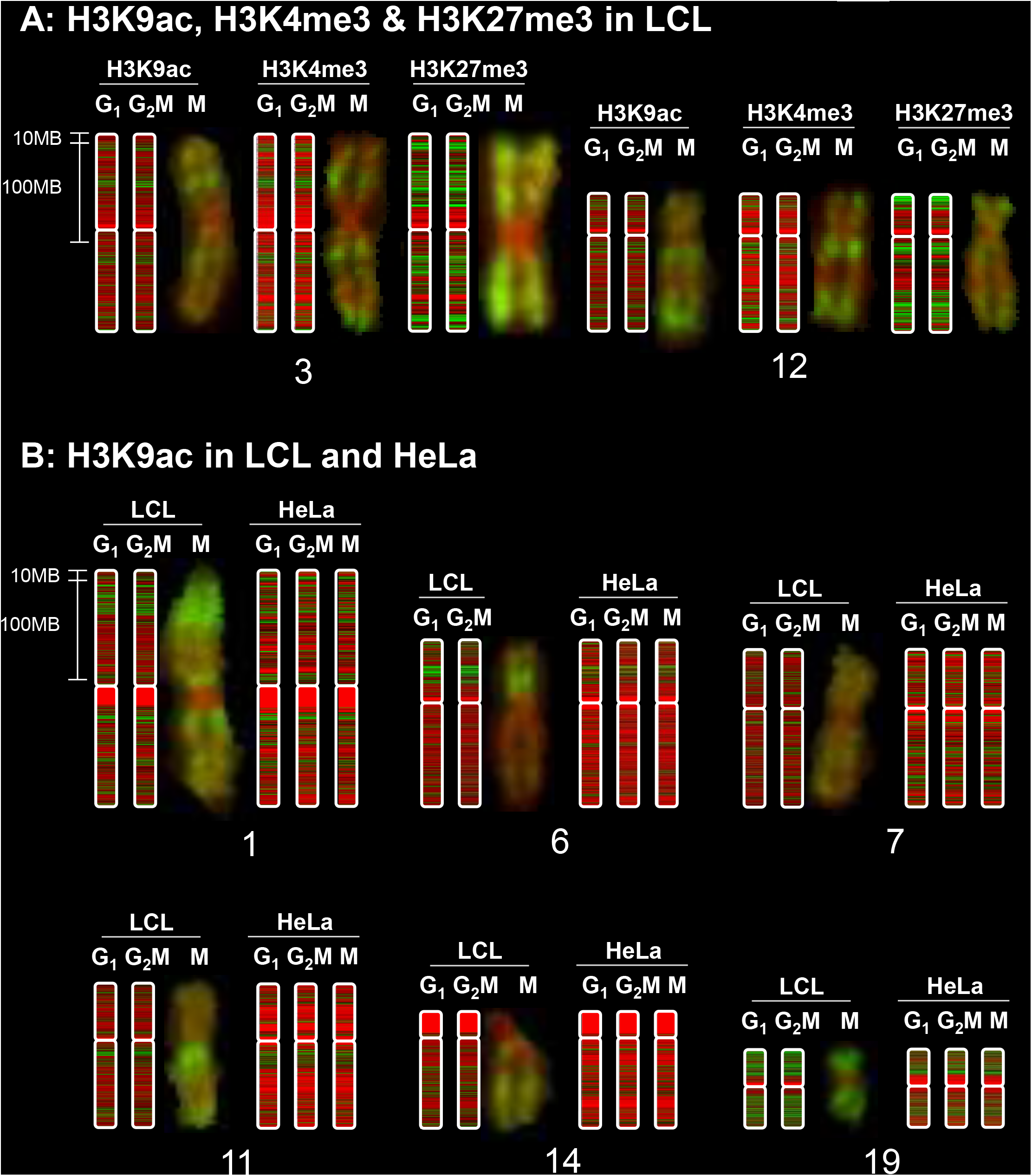
Metaphase chromosome banding patterns are conserved through the cell cycle but differ between cell types. ChIPseq data were analysed using rolling windows and displayed as blocks using a red-green colour scale in SeqMonk. A scale bar is shown, and as a rough guide, the blocks of centric heterochromatin (uniform red) at chr1q1 are 20 Mb in size. Maximum resolution (1 pixel) is approximately 1.5MB. G_1_ and G_2_M fractions were sorted by FACS (LCL) or centrifugal elutriation (HeLa). Mitotic HeLa cells were harvested by mitotic shake-off. LCL ChIP-seq karyotypes are shown aligned with metaphase chromosomes stained with antibodies to H3K9ac, H3K4me3 and H3K27me3 as described previously [32] in G_1_, G_2_M and M across selected chromosomes in LCL (A) and H3K9ac in LCL and HeLa cells (B).

The polycomb-associated modification H3K27me3 was more widely distributed across interphase chromosomes, consistent with the more diffuse banding pattern revealed by immunofluorescence microscopy of metaphase chromosomes [32]. As with the “active” modifications, immunofluorescent H3K27me3 bands detected in metaphase chromosomes were present in interphase and comprised multiple, smaller sub-bands (Figure 1A, full karyotype in supplementary figure S5).

### Chromosome sub-bands defined by histone PTM differ between cell types but are largely unchanged through the cell cycle

At microscopic (10-50Mb) resolution, there were strong similarities in banding patterns between LCL and HeLa, but clear differences in the distribution of sub-bands were visible (Figure 1B, supplementary figure S3). This is consistent with the conservation of IF banding patterns in LCL and primary lymphocytes [32,35] and with IF staining of a human x mouse somatic cell hybrid containing only human chromosome 11. The chromosome retained its characteristic banding pattern when immunostained with antibodies to H3K4me3, despite being housed in a foreign (mouse) cellular environment (Supplementary figure S6).

Visually, the distributions show only very minor differences between G_1_ and G_2_M chromosomes in LCL (Figure 1A, S3A, S4, S5) and G_1_, G_2_M and metaphase chromosomes (M) in HeLa (Figure 1B, S3B). Thus, in each cell type, every chromosome has its own distinctive and cell-type-specific pattern of bands and sub-bands. This pattern changes very little through the cell cycle, including metaphase.

These results all suggest that, at microscopic resolution (10-50Mb), histone modification patterns are dependent primarily on genomic features embedded in DNA sequence, such as gene density or frequency of repetitive elements [32]. At higher resolution (1-5Mb), cell type specific differences are apparent that may reflect cell-type determining differences in gene expression pattern.

### Correlation analysis at <1Mb resolution confirms the stability of histone modification patterns through the cell cycle

To obtain an unbiased estimate at sub-Mb resolution of genome-wide changes in histone PTM through the cell cycle, we used a rolling window analysis of our ChIP-seq data. For such analysis, the choice of window size is crucial. To examine this, we scanned our ChIP-seq data using a series of rolling windows from 1Mb to 100bp. For each window size, we calculated the correlation (Pearson R value) between cell cycle phases for H3K4me3, H3K9ac and H3K27me3. R values (R) were the plotted against window size (W). Figure 2A shows R v window size (W) for H3K4me3, K3K9ac and H3K27me3 for G_1_ vs G_2_M in LCL (left), along with H3K9ac for G_1_ vs G_2_M, G_1_ v M and G_2_M vs M in HeLa cells (right).

**Figure 2:**
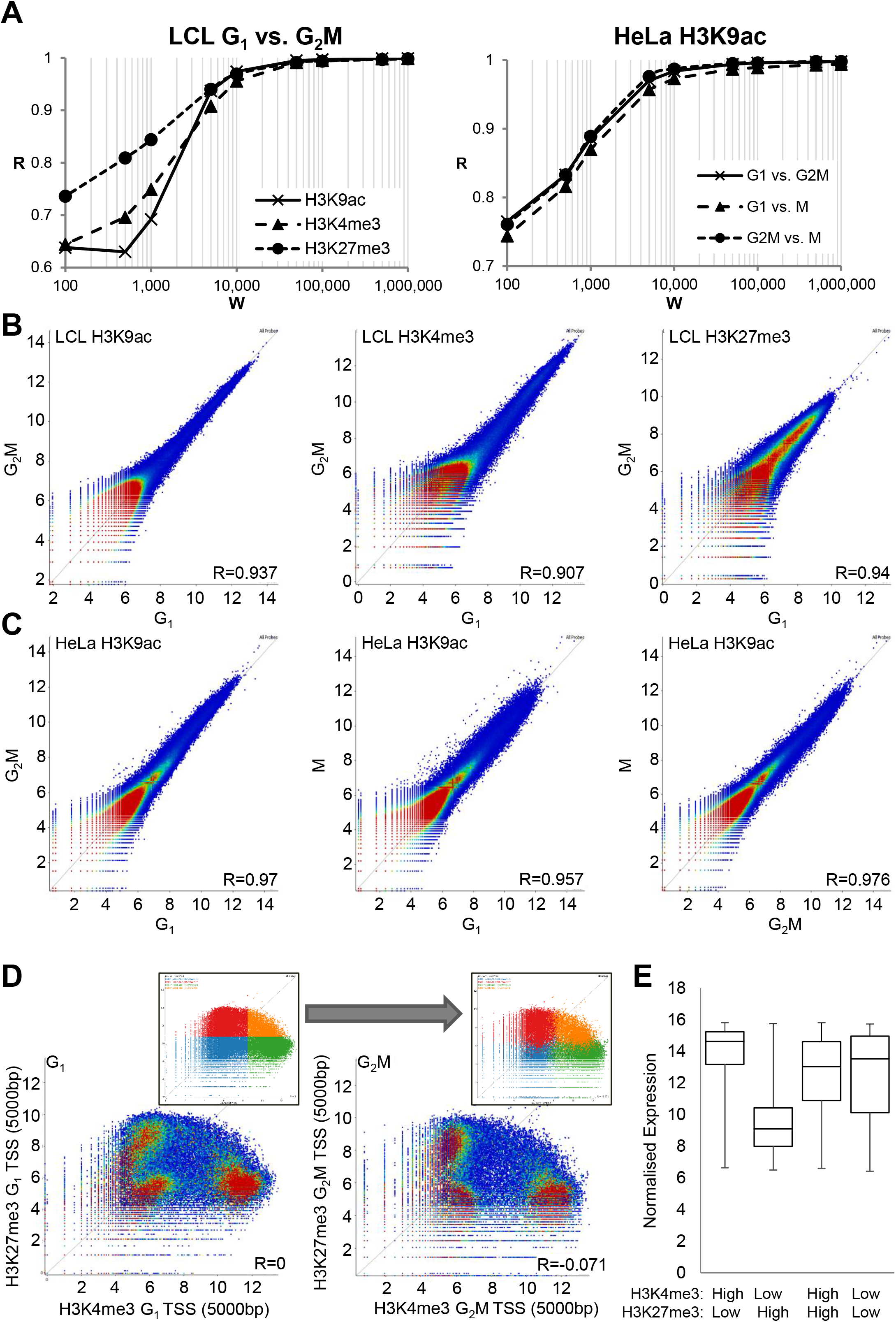
Rolling window analysis of histone modification patterns shows strong conservation through the cell cycle. (A) ChIP-seq of H3K9ac, H3K4me3 and H3K27me3 were compared in G_1_ vs. G_2_M in LCL and H3K9ac in G_1_ vs. G_2_M vs. M in HeLa cells using rolling windows of various sizes. Pearson correlation coefficient R is plotted against window size (W) for rolling windows of 100bp - 1MB. (B) 5kb windows were subsequently chosen as most informative and rolling window comparison of H3K9ac, H3K4me3 and H3K27me3 in G_1_ vs. G_2_M by dotplot is shown in FACs–sorted LCL and (C) elutriation and mitotic shake-off sorted HeLa cells. Values represent log_2_ read count normalised to total read count in a 5kb rolling window analysis with a 5kb step. Pearson R values for each correlation are indicated. (D) Using 5kb windows centred on transcription start sites (TSS), H3K4me3 was plotted against H3K27me3 in LCL in G_1_ and G_2_M. Insets show the division of the G_1_ plot into quadrants and the subsequent position of these probes in the G_2_M plot. (E) Expression levels of genes from TSS in each quadrant was measured by microarray and summarised by boxplot showing median, interquartile range, minimum and maximum.

In all cases, R was very high (>0.99) for window sizes above 50Kb. There was then an elbow at 5-10Kb below which R fell more rapidly, dropping to between 0.6 and 0.8 at a window size of 100bp. It seems that histone modification patterns are extremely well conserved through the cell cycle, even at resolutions approaching the size of a single nucleosome (200bp, figure 2A).

While the reduced value of R at window sizes below 5-10kb could be due to differences in the regional distribution (micro-banding) of histone PTM, it more likely reflects a growing number of windows containing very few or no reads at low window sizes. This inevitably leads to increased variance. In supplementary figure S7, we show that simply reducing the number of reads in the dataset progressively shifts the inflexion point (elbow) to higher values of W.

Based on these results, we selected a window size of 5Kb for further analyses. It is large enough to avoid inaccuracies caused by low integer read counts but not so large that any regions of difference between samples are lost within larger unchanging regions. Using 5Kb windows, there is a strong correlation for all modifications studied between G_1_ and G_2_M phases in LCL (R=0.907-0.94, figure 2B) and G_1_, G_2_M and M phases in HeLa (R=0.957-0.976, figure 2C).

Although genomic elements including enhancers, promoters and centric heterochromatin, are often identifiable by their exceptional levels of individual histone modifications, the identification of functionally distinct sub-categories often requires analysis of combinations of histone PTM (see [36] for examples). A particularly striking example is provided by the regions designated bivalent domains that are enriched in *both* activating and silencing histone PTM (H3K4me3 and H3K27me3 respectively) and are prominent in both mouse [37]and human [38–40] embryonic stem cells This unusual combination of PTM marks a chromatin state in which genes required for progression down specific developmental pathways, are poised to become active when the appropriate developmental signals are received.

We used scatter plots constructed from 5kb rolling windows to compare levels of H3K4me3 and H3K27me3 in G_1_ and G_2_M human LCL (figure 2D). This revealed three distinct populations, representing TSS unmarked by either modification (9,995 unique genes), TSS high in H3K4me3 and low in H3K27me3 (10,156 genes) and those low in H3K4me3 and high in H3K27me3 (8,315 genes). In these differentiated cells only a small proportion of genes (2,825 genes) were present in bivalent domains, relatively highly enriched in both marks. This pattern was consistent between cell cycle phases and TSS retained their position in the distribution from G_1_ to G_2_M (figure 2D, insets). Expression levels from genes in each population were determined by microarray from asynchronous cells (figure 2E). As might be expected, expression was highest from TSS with high H3K4me3 and low H3K27me3 and this group was enriched in genes involved in housekeeping processes such as mitochondria (P=2.5×10^−18^), ribonucleoprotein complex (P=9.6×10^−14^) and RNA processing (P=1.6×10^−9^).

Expression was lowest from TSS with high H3K27me3 and low H3K4me3 and these genes were enriched in cell-type specific genes which would not be expected to be expressed in LCLs such as epidermis development (P=8.5×10^−8^), neuron differentiation (P=3.3×10^−5^) and epithelial cell differentiation (P=2.4×10^−5^). Genes with high levels of H3K4me3 and H3K27me3 or low levels of both marks at their TSS showed a broader range of expression, suggesting these two marks were not sufficient to define their transcriptional status. Despite LCLs being differentiated cells, Hox genes were enriched within the H3K4me3 high, H3K27me3 high “bivalent” group of TSS (fold enrichment 3.1, P=0.003). It has been shown that in human ES cells, the level of H3K4me3 at bivalent promoters varies through the cell cycle, with some genes showing elevated H3K4me3 exclusively at mitosis. Intriguingly, these genes showed the strongest up-regulation after induction of differentiation. Further, in differentiated cells, levels of H3K4me3 at bivalent domains became stable through the cell cycle [40], a finding consistent with our results in LCL. In differentiated cells, it was shown that the writers of active modifications generally dissociate from mitotic chromatin while writers of silencing marks, including members of the polycomb complex are retained [21],

### Differences in H3K9ac distribution between cell types reflect cell-type-specific transcription

When H3K9ac modification patterns were compared between HeLa and LCL, correlation was lower than between cell cycle phases, though still high overall (G_1_ P=0.781, figure 3A, G_2_M P=0.808, not shown). A small number (~6%) of windows showed greater difference, with relatively high modification levels in HeLa or LCL, indicative of cell-specific PTM levels. To explore this further, the 3% of windows exhibiting the highest differential enrichment in H3K9ac in HeLa relative to LCL and *vice-versa,* were selected and the associated genes extracted. These outlying windows were relatively gene dense (27 genes/Mb for the LCL>HeLa group, 57 genes/Mb for the HeLa>LCL group, compared with 14.1 genes/Mb for the whole genome). Expression levels from genes within these regions were analysed by microarray, revealing that H3K9ac enrichment correlated with gene expression: those probes most enriched in H3K9ac in LCL compared to HeLa contained genes more highly expressed in LCL and vice versa (figure 3B). Ontology analysis revealed that windows more enriched in H3K9ac in LCL than HeLa cells contained genes involved in B-cell receptor signalling pathway (fold enrichment (FE) 5.9, P=3.6×10^−15^), adaptive immune response (FE 2.9, P=9.9×10^−9^) and MHC class II protein complex (FE 5.3, P=4.3×10^−5^). In contrast, regions more H3K9ac-enriched in HeLa contained genes involved in pathways in cancer (FE 1.5, P=9.9×10^−8^), including oncogenes such as ERBB2, WNT5A and BRAF.

**Figure 3:**
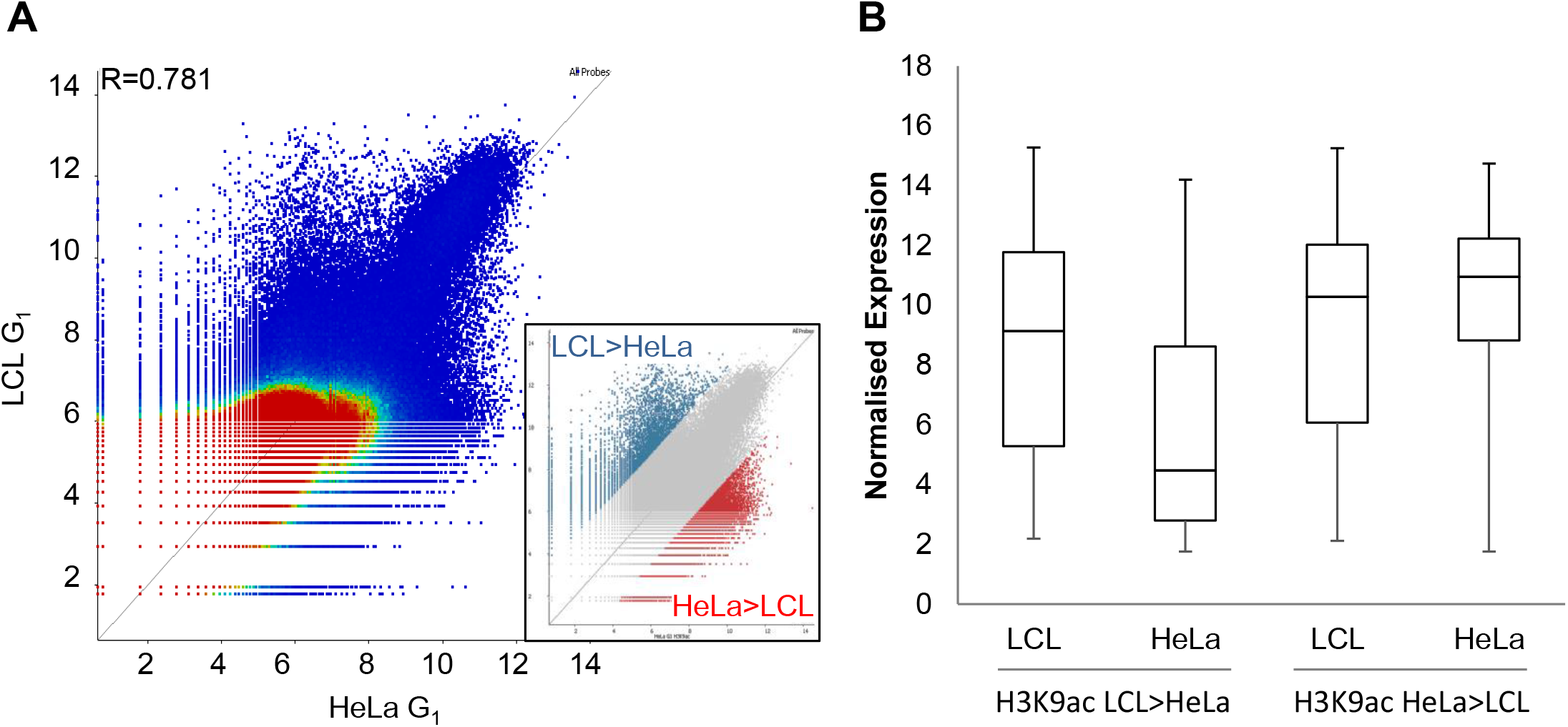
Differences in H3K9ac pattern between LCL and HeLa reflect cell-type-specific transcription. (A) Dotplot comparison of H3K9ac in 5kb windows between LCL and HeLa in G_1_. The outlying 3% on each side of the distribution are highlighted (inset). (B) The expression of genes contained within these outlying regions was measured by microarray in LCL and HeLa.

### Histone modification levels at Topologically Associated Domains do not change through mitosis

As a first step towards linking sub-microscopic histone PTM banding with genomic function, we examined histone modification levels within topologically associated domains, TADs [15,16]. These domains range in size from about 0.5 to 2Mb, with a median size of around 0.9Mb [15]. They show a characteristic distribution of histone modifications, including H3K4me3 and H3K27me3 [16],with boundary elements appearing to constrain the spread of heterochromatin [15]. TAD structure is lost as chromosomes condense during mitosis and re-emerges during early G_1_ [18]. We quantified ChIP-seq read counts within TADs using TAD coordinates defined by Dixon et al. [15] in IMR90 human fibroblasts. We compared levels of H3K4me3, H3K9ac and H3K27me3 in G_1_ vs G_2_M in LCL, and H3K9ac in G_1_ vs G_2_M vs M in HeLa cells. We found that histone modification levels within TADs were strongly conserved from G_1_ into G_2_M and M in both LCL (R=0.979 −0.994, figure 4A) and HeLa (R=0.981 −0.996, figure 4B). As shown in figure 4C, distribution of histone PTM across TADs, particularly that of H3K27me3, appears to be confined by domain boundaries and the higher resolution arrowhead domains described by Rao et al. [16]. However there was no reduction, in signal or redistribution from G_1_ into G_2_M and M, despite the disappearance of detectable TADs in metaphase [18].

**Figure 4:**
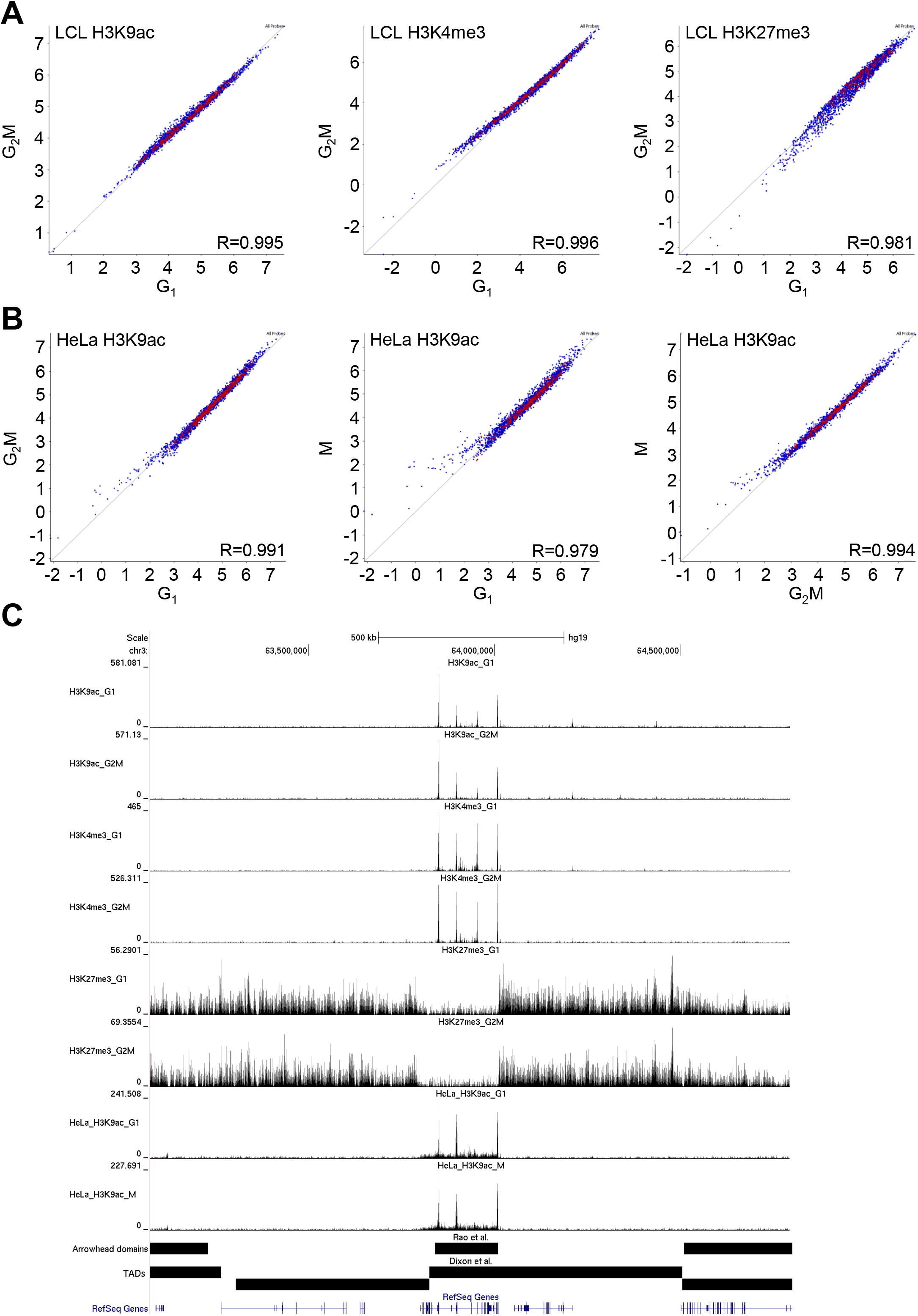
Histone modification levels are conserved within topological domains throughout the cell cycle. (A) Comparison of H3K9ac, H3K4me3 and H3K27me3 in G_1_ vs. G_2_M in FACs–sorted LCL and (B) H3K9ac in HeLa in G_1_ vs. G_2_M, G_1_ vs. M and G_2_M v M (elutriated / MSO). Topological domain coordinates from Dixon et al. [15] were used to define probes. Values represent log_2_ read count normalised to total read count, corrected for probe size. Pearson R values are given for each correlation. (C) A genome browser screenshot of cell-cycle specific ChIP-seq data on chromosome 3. All tracks are from LCL except where indicated. Topological domains [15] and higher-resolution arrowhead domain [16] are shown.

TADs contain more than a single gene, and these can be active, inactive or a mixture of both. Similarly, most TADs contain a mixture of “activating” (H3K9ac, H3K4me3) and “silencing” (H3K27me3) histone modifications, the former distributed as sharp peaks (<5Kb), marking DNA elements such as promoters or enhancers and the latter as broader genomic regions, sometimes incorporating several control elements or coding regions. This is all consistent with previous data [4,36,41], but what is remarkable is the strict, general conservation of these distributions through the cell cycle, despite the structural reorganization involved. Our results raise the interesting possibility that a conserved pattern of histone modification facilitates the reassembly of TADs in a way that preserves their distinctive transcriptional properties.

### Gene function and expression timing modulate histone modification at specific transcription start sites

*We* next focussed on the regulatory regions around transcription start sites (TSS). The average distribution of H3K4me3 across TSS showed a characteristic bimodal profile around TSS in LCL (Figure 5A), with the expected nucleosome-free region [42] and the highest modification levels at the +1 nucleosome, decreasing over subsequent nucleosomes to basal levels by approximately 2000bp into the gene body. The average distribution did not vary significantly between G_1_ and G_2_M. We saw very similar average distributions around TSS for a second modification, H3K9ac (Figure 5C). Using TSS windows from −500bp to +750bp, approximately incorporating the first nucleosome upstream of the nucleosome free region (−1) and the first three downstream nucleosomes (+1 to +3), we showed that H3K4me3 and H39ac levels at TSS correlated very strongly between G_1_ and G_2_M (R=0.964 and 0.967 respectively, figure 5B & 5D).

**Figure 5:**
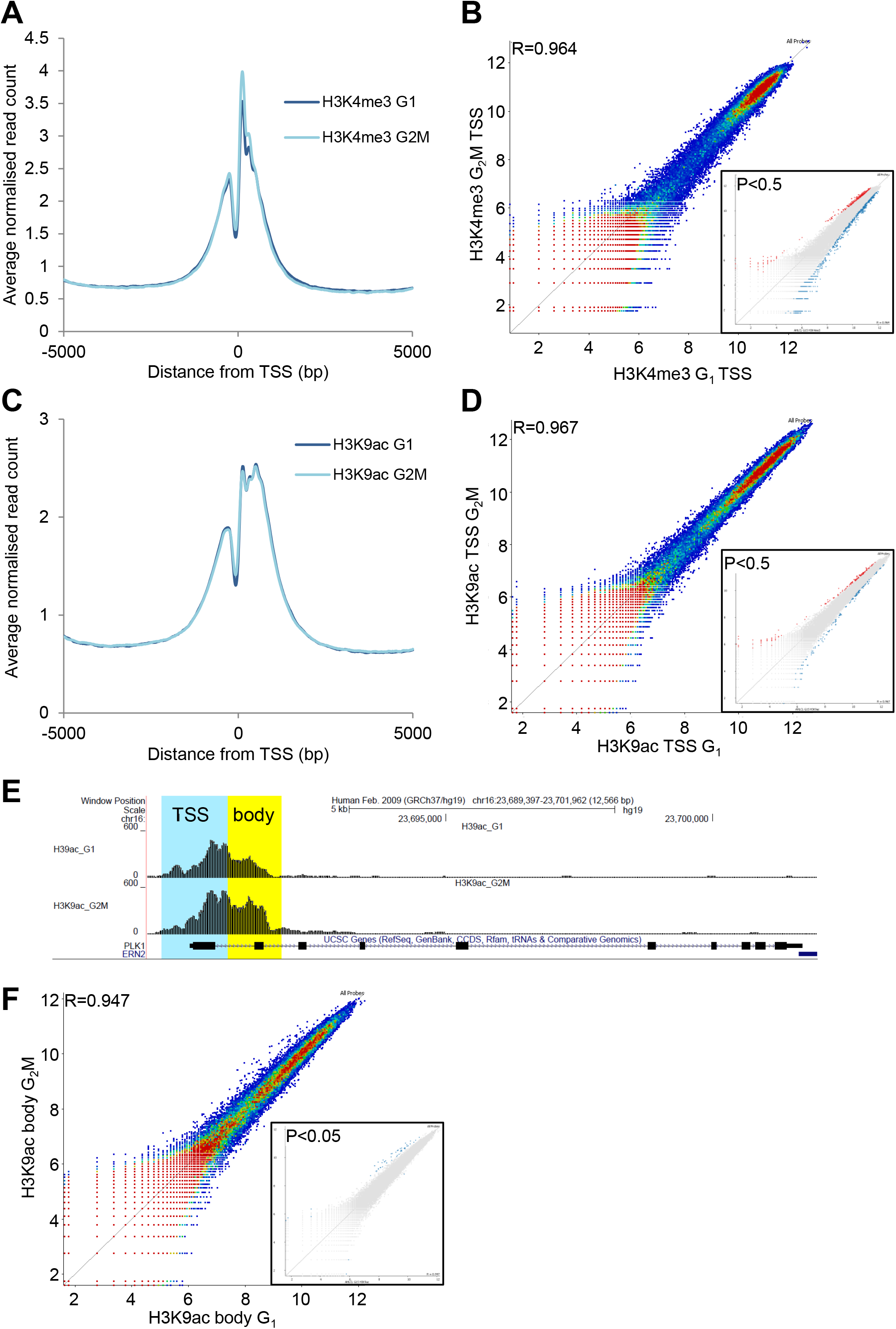
H3K4me3 and H3K9ac enrichment at transcription start sites is conserved in LCL, with outliers representing specific gene function. (A, C) Average distribution (average read count normalised to total read count) across a 10kb window centred on all transcription start sites (TSS) for H3K4me3 (A) and H3K9ac (C) in G_1_ and G_2_M in LCL. (B, D) Modification levels at the proximal TSS (−500 to +750bp) are compared by dotplot for H3K4me3 (B) and H3K9ac (D). Outliers were selected by intensity difference (P<0.5, insets) for ontology analysis. (E) Extension of the H3K9ac signal further into the gene body (+750 to +1750bp) was observed for a few genes with function in mitosis, e.g. PLK1. (F) Comparison of H3K9ac enrichment in this region is shown and outliers highlighted (P<0.05, inset).

Despite the strong correlations shown by these scatter plots, there are outliers towards the edges of the distributions (Figure 5). We asked whether these have any functional significance, perhaps representing distinct classes of genes. Intensity difference (P<0.5) was used to identify outliers (Figure 5B & 5D, inset panels). These genes were subjected to ontology analysis using DAVID. Strikingly, gene populations selected for high levels of either H3K4me3 or H3K9ac in G_2_M relative to G_1_, were consistently and strongly enriched in GO terms related to mitosis (supplementary table S1, P=4×10^−11^ −0.004, fold enrichment (FE) 2.3-10.4). When TSS significantly enriched in H3K9ac in G_1_ or G_2_M were highlighted on the H3K4me3 scatterplot (supplementary figure S8A), they aligned to the same side of correlation and *vice-versa* (supplementary figure S8B), showing that at higher, i.e. more accurate, levels of modification, the two PTM are closely correlated.

We also observed that at some TSS with H3K9ac enrichment in G_2_M, modification levels extended further into the gene body in G_2_M than in G_1_ (e.g.PLK1, Figure 5E). We therefore correlated gene body (+750bp to +1750bp) acetylation levels (figure 5F) in G_1_ and G_2_M. Correlation was again strong with a small group of outliers, which were statistically more significant (P<0.05) than those for TSS and were almost entirely enriched in G_2_M relative to G_1_ (figure 5F, inset). This small group of genes was extremely strongly enriched in mitotic terms (supplementary table S1): genes annotated as involved in mitotic cell cycle comprised 10 of 26 genes enriched in the gene body in G_2_M, representing a fold enrichment of 19.2 (P=3.0×10^−10^).

It may be that increased H3K9ac in G_2_M is due to transcription, late in the cell cycle, of genes whose products are required for mitosis. To test this, we used centrifugal elutriation to isolate 4 fractions of lymphoblastoid cells at sequential points through the cell cycle (figure 6A). We then carried out RT-PCR using primers specific for the pre-mRNA of genes previously shown to have particularly high H3K9ac in G_2_M, spreading across the TSS and into the gene body. The results are shown in Figure 6B. Of 7 genes with elevated H3K9ac in G_2_M, transcript levels were always highest in fractions 3 and 4, consisting predominantly of G_2_M cells. Conversely, a single gene (NBN) with comparably elevated H3K9ac in the gene body in G_1_ cells and genes with elevated H3K9ac or H3K4me3 at the TSS in G_1_ or G_2_M, showed slightly elevated transcription in fractions 1 and 2 (Figure 6B), containing a majority of G_1_ cells (Figure 6B).

**Figure 6:**
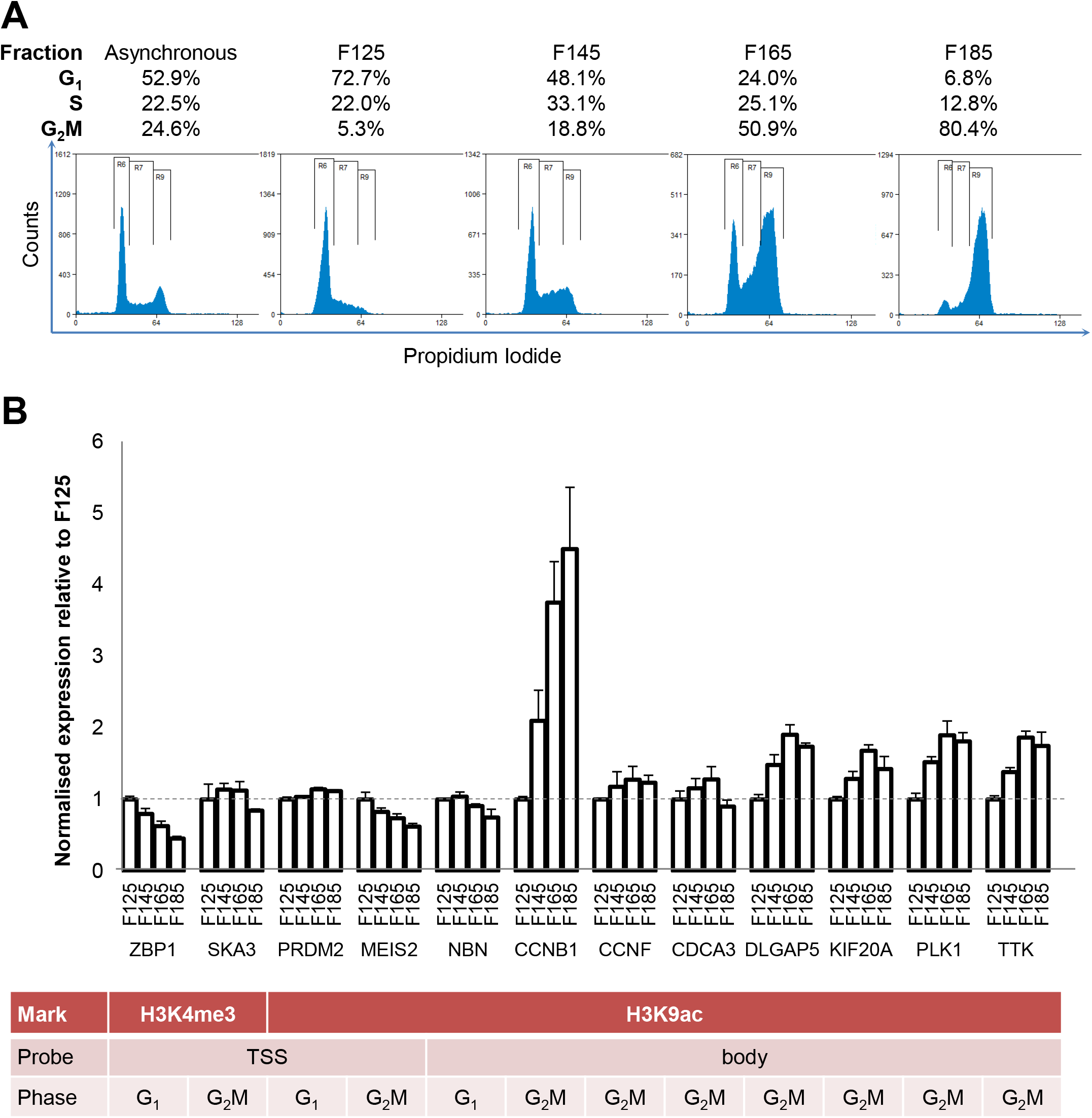
H3K9ac gene body enrichment in G_2_M is associated with ongoing transcription. (A) LCL were fractionated by centrifugal elutriation. Elutriated fractions were fixed and stained with propidium iodide to determine cell cycle stage. The proportion of cells in G_1_, S and G_2_M for each elutriated fraction is shown above each plot. (B) Cell-cycle specific expression of genes with significant TSS enrichment was measured by RT-qPCR with primers to nascent pre-mRNA, normalised to the housekeepers β-actin, beta-2-microglobulin, RPLI3A and HPRT. Genes were enriched in H3K4me3 at the TSS in G_1_ (ZBP1) or G_2_M (SKA3), H3K9ac at the TSS in G_1_ (PRDM2) or G_2_M (MEIS2) or in the gene body (+750 - +1750) in G_1_ (NBN) or G_2_M (all remaining genes).

H3K9ac enrichment at TSS was also strongly conserved through the cell cycle, including metaphase, in HeLa cells, (G_1_ vs G_2_M, R=0.978; G_1_ vs M, R=0.967; G_2_M vs M, R=0.975, figure7). When minor outliers at the edges of the correlation were identified (P=0.5, figure 7B, inset panels), mitotic terms were again over-represented in TSS enriched in G_2_M and M relative to G_1_ (P=0.004 and P=0.003, FE=3.0 and 1.6 respectively, figure 7, supplementary Table S2).

**Figure 7:**
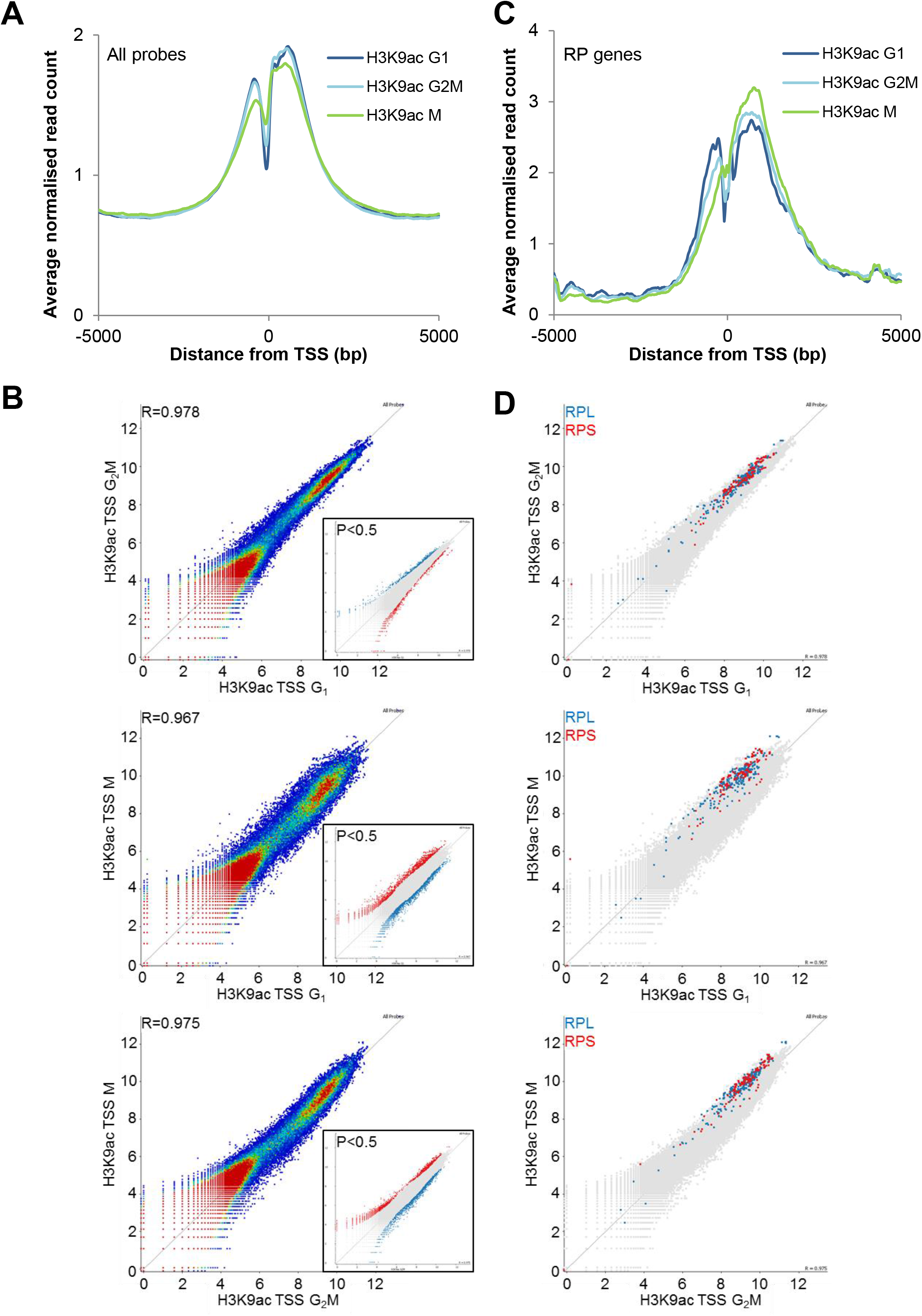
Transcription start sites of ribosomal protein genes are enriched in H3K9ac in mitosis. (A) HeLa cells were sorted by centrifugal elutriation (G_1_, G_2_M) and mitotic shakeoff (M). Average distribution (average read count normalised to total read count) across a 10kb window centred on all transcription start sites (TSS) is shown. (B) ChIP-seq signal at TSS (−500bp to +750bp) is compared between cell cycle phases. Significantly enriched TSS (P<0.5) were identified by intensity difference as shown in the inset panels. (C) The average H3K9ac distribution across the TSS of ribosomal protein (RP) genes (large and small, excluding mitochondrial) is shown. (D) ChIP-seq signal at TSS of ribosomal protein genes is compared between cell cycle phases. RPL genes are shown in blue and RPS genes in red.

Thus, increased histone acetylation is closely linked to the selective expression of a group of genes necessary for mitosis at the appropriate stage of the cell cycle. The change in acetylation occurs in the same cell cycle phase (G_2_) as the increase in transcription, suggesting that it is part of the transcription process rather than a pre-determining marker of selected genes. A more detailed temporal analysis of the two events will be necessary to resolve this.

In HeLa, but not LCL, translational terms, particularly ribosomal protein genes, were strongly enriched in H3K9ac in G_2_M relative to G_1_ (P=1.1×10^−21,^ FE=18). Surprisingly, ribosomal protein genes were further enriched in H3K9ac in M phase relative to G_2_M (supplementary Table S3), suggesting that differential marking is maintained through mitosis. Large and small ribosomal genes (28 of each) were all enriched in M relative to G_2_M (P=2.2×10^−14^ and 1.4×10^−13^ FE= 5.7 and 5.4 respectively, Supplementary Table S2). The position of RPL and RPS genes in the correlation scatterplots are shown in figure 7D, showing a strong bias towards enrichment in mitosis. M phase enrichment is also accompanied by a change in the average modification profile with a reduction in H3K9ac enrichment at the −1 and +1 nucleosomes from G_1_ to G_2_M to M and higher enrichment further into the gene body (Figure 7C), perhaps reflecting ongoing transcription of RP genes.

The H3K9ac around the TSS of RP genes extends further into the gene body, at all stages of the cell cycle, than TSS in general (all probes). In view of the results with mitosis-related genes preferentially transcribed in G_2_, this suggests that RP genes may be transcribed throughout the cell cycle, including through mitosis. Whether transcription itself persists or not, the finding that levels of H3K9ac at RP TSS are higher in metaphase than any other stage of the cell cycle, suggests a chromatin mark that helps maintain the continuity of RP gene transcription through mitosis. Continued transcription is also consistent with our failure to detect any increase in RP transcript levels (compared to genes in general) as HeLa cells emerged from mitosis into G_1_.

## DISCUSSION

We have asked how the genomic distribution of three histone modifications closely implicated in both gene expression and chromatin structure, H3K9ac, H3K4me3 and H3K27me3, differs between cell types and through the cell cycle. We have used ChIP-seq to define the distribution of these modifications in G_1_, G_2_M and M phase cells from two cell types (HeLa and lymphoblastoid cell lines, LCL) at various levels from 10-50Mb chromosome domains (bands) down to single gene promoters and Transcription Start Sites (TSS).

We find that the 10-50Mb bands previously shown to be relatively rich in H3K9ac, H3K4me3 and/or H3K27me3 by immunofluorescence microscopy of human metaphase chromosomes [32], are all detectable by ChIP-seq. in interphase (G_1_, G_2_) and metaphase cells. At all cell-cycle phases and in both cell types, they are seen to be assemblies of smaller sub-bands (1-2Mb), below the limit of detection by conventional microscopy.

The same large bands across each chromosome are present in both HeLa and LCL. In contrast, at 1-2Mb, sub-band resolution, clear differences between the two cell types are visible. From this, it is reasonable to conclude that the 10-50Mb IIF bands are specified by genome sequence whereas the small sub-bands are linked to the pattern of gene expression characteristic of each cell type.

Remarkably, at all levels of resolution, there was little or no change through the cell cycle. Thus, at 1-2Mb resolution and below, each chromosome has a distinctive and celltype-specific banding pattern that is essentially the same in G_1_, G_2_ and metaphase.

### Histone PTM and maintenance of cellular identity through the cell cycle

Chromatin controls transcription in eukaryotes at multiple levels, from nucleosome packaging (repeat length of around 200bp), to chromatin loops (exemplified by TADs, between 0.2 – 1.5 Mb) and various sub-nuclear structures visible at the light microscope level, such as blocks of centric heterochromatin, Polycomb bodies and nuclear speckles [43–46]. Structures can be built by chromatin looping mediated by protein-protein and protein-DNA binding [47,48] and BY the spontaneous collapse of large chromatin domains at high concentration, leading to phase transitions and highly condensed blocks of chromatin [6,49]. Phase separation can be influenced by DNA sequence [50], by overall levels of histone acetylation and bromodomain binding proteins [14] and by DNA methylation [51], the latter two providing potentially mechanistic links between higher-order chromatin structures common epigenetic marks. These structures influence transcription in various ways; they can restrict access of TFs and other functional proteins to their binding sites, and bring regulatory DNA elements (such as enhancers and promoters) into spatial proximity [52]. A key point for the present discussion is that these structures are all dynamic [53] and subject to disruption through the cell cycle; TADs and higher order structures are disrupted, and often disappear altogether, through mitosis [53,54]. How then are cell-type-specific patterns of gene expression to be maintained?

To explore in more detail the link between histone PTM levels and transcription through the cell cycle, we scanned histone PTM across LCL and HeLa genomes using rolling 5Kb windows. On scatter plots, a consistently close correlation was found when different cell cycle phases were compared (R values between 0.91 and 0.98 for all three modifications). When the same procedure was used to compare H3K9ac at equivalent cell cycle phases in HeLa and LCL, the correlation, though still present, was much lower (R=0.78). Further, outlying regions on either the LCL or HeLa sides of the distribution, were enriched in genes likely to be preferentially expressed in either HeLa or LCL, i.e. cell-type-specific genes. These results are consistent with the proposition that variation in PTM distribution at 1-2Mb and below, reflects the distinctive patterns of transcription, or transcriptional potential, that characterize the two cell types, LCL and HeLa.

We find that histone modifications associated with TADs [15,16] show cell-specific differences and are conserved through the cell cycle including, in the case of H3K9ac and HeLa cells, passage through metaphase. This is significant in that TADs are undetectable by chromatin conformation capture in metaphase HeLa cells, but reappear a few hours into G_1_ [18].

Levels of histone PTM at Transcription Start Sites, are even more highly correlated across the cell cycle than levels within unselected windows (R=0.964 and 0.978). It seems that modification levels at TSS are almost universally invariant through the cell cycle. However, we identified a small group of genes with relatively high levels of H3K9ac and H3K4me3 across their TSS in G_2_M. Ontology analysis showed this group to be highly enriched in genes whose products are required for mitosis. We showed that primary transcript levels of selected members of this gene family increased as cells progressed into G_2_, but levelled off towards the end of this phase, suggesting that the increase in activating modifications reflects on-going transcription. This conclusion is consistent with the finding that the increase in G_2_M of H3K9ac in the *gene body* is even more marked than the increase at TSS.

Our results point to the existence of a homeostatic mechanism that serves to maintain levels of certain histone PTM at specific locations, through the cell cycle. This is a challenging proposition, not least because histone acetylation and methylation are both dynamic processes maintained by the balanced action of appropriately targeted modifying and de-modifying enzyme complexes [1]. Thus, the unchanging distribution and relative levels of H3K9ac, H3K4me3 and H3K27me3 through the cell cycle, requires an unchanging balance of specifically targeted enzyme activities. The mechanism must permit the persistence of local levels of histone PTM when new chromatin is assembled during S-phase. Further, modifications that show very little cell-cycle-dependent change, such as H3K9ac, must be managed alongside those that change rapidly at specific cell cycle stages, such as H3S10ph in G_2_ [24–26]. These considerations make it unlikely that levels of modifications such as H3K9ac are conserved solely through adoption of a chromatin configuration that excludes the relevant enzymes. Nor can the maintenance of patterns of modification be attributed solely to on-going transcription, important though this is in the particular case of mitotic genes expressed in G_2_ (see above). Firstly, transcription of most genes is suppressed through mitosis [55] though there are interesting exceptions [56,57].

Whatever the mechanism may prove to be, previous data support the existence of a mechanism conserving levels of at least some histone modifications. For example, treatment of cultured cells with HDAC inhibitors (HDACi) such as sodium butyrate, causes little or no increase in histone acetylation across most genes tested, despite global histone hyperacetylation [58,59]. Further, within 60 minutes of exposure to HDACi, we noted decreased transcription of genes encoding essential components of all known HAT complexes, a change suggestive of a homeostatic process aimed at conserving levels of histone acetylation. The initially surprising inability of HDACi to induce hyperacetylation at most gene promoters is consistent with the limited transcriptional response to HDACi, and the fact that genes that do respond can be up- or down-regulated [60–62].

### Marking selected genes through mitosis

There is a general reduction in histone acetylation as cells move into mitosis, driven largely by the actions of histone deacetylases [63]. Moreover, turnover of various histone acetates is diminished in metaphase HeLa cells, as measured by the minimal hyperacetylation induced by treatment with HDAC inhibitors such as sodium butyrate [64]. Despite this overall change in turnover rate, we see very little change in *the relative* levels of modification of the regions studied here. With few exceptions, they all respond in the same way.

It has been proposed that patterns of expression of at least some genes are conserved through mitosis by the retention of selected transcription factors at the promoter regions [19,20,65,66].

The hypothesis has been termed bookmarking, and recent work has shown that it can involve not only TFs, but enzymes such as the p300 complex [67] and poly [ADP-ribose] polymerase I [68], nucleosomes [69] and histone PTM [70], see [71] for a recent review. These findings are all compatible with the multiple levels of control that we now know to be involved in transcriptional regulation in eukaryotes.

Among the small number of genes showing disproportionate changes in acetylation through metaphase, levels of H3K9ac were consistently elevated at Transcription Start Sites of ribosomal protein (RP) genes in metaphase HeLa cells. It is unclear why these very highly expressed housekeeping genes [72] should respond differently to other genes or genomic domains, but it is significant that all the RP genes we have tested respond in the same way. The balanced expression of the 80 members of the RP gene family is essential for efficient ribosome biogenesis [73]. Thus, although the functional significance of the increased H3K9ac at RP gene promoters remains unknown, it is of interest that all members of the family tested showed the same characteristic marking through metaphase. Again, the observation is consistent with a mechanism involving specific histone modifications, that maintains the particular functional characteristics of this gene family through mitosis. It is interesting that the rRNA genes, despite being expressed in all cells, are marked through mitosis by the transcription factor Runx2 in a lineage-specific manner [74].

## CONCLUSIONS

Genome-wide analysis by ChIP-seq, shows histone modifications H3K4me3, H3K9ac and H3K27me3 to be relatively enriched over chromosomal domains of 10-50Mb. These domains do not differ detectably between cell types, do not vary through the cell cycle and correspond to bands detectable by immunofluorescence microscopy of metaphase chromosome spreads. The relatively high levels of both activating and silencing histone PTM in these regions are likely to be by-products of their high gene density.

By ChIP-seq, these regions are made up of clearly defined sub-regions of 2-5MB or less. These sub-regions differ between cell types but are invariant through the cell cycle.

Analysis of individual promoter regions confirmed a general lack of variation through the cell cycle within each cell type, the only exceptions being genes whose transcription was up-regulated in G_2_in order to generate proteins required for passage through mitosis.

Our results suggest that H3K4me3, H3K9ac and H3K27me3 are components of a homeostatic mechanism by which chromatin states responsible for cell-type-specific patterns of gene expression are set and maintained through the cell cycle.

## METHODS

### Cell culture

The AH-LCL lymphoblastoid cell line was established by Rowe and colleagues [75] and was cultured in RPMI 1640 medium, 10% foetal bovine serum, supplemented with L-glutamine (2mM) and penicillin/streptomycin (all reagents from Life Technologies) at 37°C, 5% CO_2_). HeLa cells were cultured in DMEM medium supplemented with 10% foetal bovine serum (Sigma), supplemented with L-glutamine (2mM) and penicillin/streptomycin (Life Technologies). The human/mouse hybrid cell line GM11941, containing human chromosome 11 was purchased from the Coriell Institute and cultured in RPMI 1640 medium, 10% foetal bovine serum, supplemented with L-glutamine (2mM) and penicillin/streptomycin. All cell lines were regularly tested for mycoplasma.

### Cell Sorting Methods

#### Flow cytometry

1×10^8^ LCL yielded approximately 6×10^6^ G_2_M cells from a 4 hour sort. Cells were treated with 0.1μg/ml colcemid for 4 hours to enrich the mitotic fraction. Cells were then harvested by centrifugation and washed three times in PBS. Ice cold acetone was added dropwise to ~1×10^7^ cells/ml on a very slow vortex before freezing to −20°C for at least 2 hours. Cells were then centrifuged at 30g for 10mins, 4°C (MSE chillspin), acetone was carefully removed from the pellet and ice cold PBS was added dropwise to ~1×10^7^ cells/ml. The cells were centrifuged and resuspended in PBS at 2 x 10^7^ cells/ml. Propidium iodide was added to a concentration of 10μg/ml and incubated for 30 minutes at 4°C. Cells were sorted into G_1_ and G_2_M fraction based on Propidium iodide fluorescence using a MoFlo cell sorter (DakoCytomation).

#### Centrifugal elutriation

Centrifugal elutriation was carried out using a JE-5.0 elutriating centrifuge (Beckman Coulter) as described by Banfalvi [76]. 3×10^8^ LCL were harvested by centrifugation and resuspended in 10ml elutriation buffer (5mM EDTA, 1% bovine serum albumin in 0.75x PBS); HeLa cells were trypsinised, washed in culture medium and 3×10^8^ cells resuspended in 10ml of elutriation. A single cell suspension was achieved by passage through an 18G needle and cells were loaded into the elutriation chamber. For LCL, loading was at a flow rate of 11ml/min and a rotor speed of 1,800 rpm. For HeLa cells, the optimal G_1_ fractions was achieved loading at 15ml/min, 1800rpm and the purest G_2_M fraction at 11ml/min, 1300 rpm. Cell cycle fractions were eluted by gradually (1ml/min/min) increasing the flow rate and collecting 200ml fractions at ~3ml/min increments. Eluted cells were centrifuged and acetone-fixed as above for ChIP-seq. A sample of each fraction was fixed in 70% ethanol and cell cycle stage determined by propidium iodide staining and flow cytometry.

#### Mitotic shake-off

HeLa cells were grown on 15cm culture plates (Corning), ensuring they were in log phase and no more than 70% confluent at time of shake off. Plates were shaken on an orbital shaker at 200 rpm for 1 minute. Medium containing shaken-off cells was collected and cells pelleted by centrifugation. The first shake-off was discarded as it contained any dead cells and debris. Several subsequent shake-offs were carried out for each set of culture plates at intervals of 1 hour. Mitotic samples were collected for ChIP-seq or replated for synchronous progression into G_1_. After the desired culture time, replated cells were detached by trypsinisation and washed in growth medium followed by PBS. Cells pellets were frozen prior to extraction for western blotting. Cells were acetone-fixed as above for ChIP-seq. A sample of each fraction was fixed in 70% ethanol and cell cycle stage determined by propidium iodide staining and flow cytometry.

### Antibodies

Rabbit polyclonal antisera to H3K9ac (R607) and H3K4me3 (R612) were raised in-house by immunisation with synthetic peptides conjugated to ovalbumin as previously described [8 (White DA et al., Methods 1999, 19:417–245). Antibody specificities was assayed by inhibition ELISA and checked by Western blotting. The antibody to H3K27me3 was from Millipore (07-449).

### Chromatin Immunoprecipitation-sequencing

Immunoprecipitation of native chromatin was performed based on the method described previously [77], with modifications but using acetone-fixed cells as described above. Acetone fixation permeabilises cell and nuclear membranes, removing the need to isolate nuclei prior to micrococcal nuclease digestion. Instead, acetone-fixed cells were washed twice in cold PBS and resuspended in digestion buffer (0.32M sucrose, 50mM Tris/HCl (pH7.4), 4mM MgCl_2_, 1mM CaCl_2_, 5mM Na Butyrate, 0.1mM PMSF) at 8×10^7^ cells/ml (G_1_) or 4×10^7^ cells/ml (G_2_M). Micrococcal nuclease was added to 25units/ml and chromatin digested at 37°C for 5 minutes. The reaction was stopped by the addition of EDTA to 5mM. Chromatin was pre-cleared by incubation with protein A sepharose beads and incubated overnight with antibodies to H3K9ac, H3K4me3 and H3K27me3. Antibody-bound material was isolated on Protein A-Sepharose beads (Invitrogen, UK) and DNA from antibody-bound and input chromatin was purified by PCR purification kit (Qiagen). Sequencing libraries were prepared from 100ng DNA per sample using the NEBNext Library preparation kit for Illumina (NEB). Samples were barcoded using the system described by Bronner et al. [78]. Fragments were amplified with 12–18 cycles using adaptor specific primers (Illumina); fragments ranging between 300 and 500⍰bp in size were gel-purified before cluster generation and sequencing. Sequencing was carried out at the Babraham Institute, Cambridge on an Illumina Genome Analyzer GAIIX using Cluster Generation v4 chemistries and Sequencing by Synthesis Kit v4. Data collection was performed using Sequencing Control Software v2.6. Real-time Analysis (RTA) 1.5-1.8 were used for base calling. ChIP-seq data was analysed using SeqMonk (Babraham Institute, http://www.bioinformatics.babraham.ac.uk/projects/seqmonk/).

### Immunofluorescence

Metaphase chromosome spreads were prepared and immunostained as previously described (Terrenoire et al. 2010). Briefly, cells were treated for two hours with colcemid at 0.1 μg/ml. Cells were washed twice with cold PBS and swollen in 75mM KCl at 2×10^5^ cells/ml at room temperature for 10min. 200μl aliquots of the swollen cell suspension were spun onto glass slides at 1800 rpm for 10min in a Shandon Cytospin 4. Slides were immersed for 10min. at room temperature in KCM buffer (120mM KCl, 20mM NaCl, 10mM Tris/HCl pH 8.0, 0.5mM EDTA, 0.1% Triton X-100). Immunolabelling was carried out for 1 hour at 4°C with antisera diluted 200-400 fold in KCM supplemented with 1-1.5%. BSA. The secondary antibody was FITC-conjugated goat anti-rabbit immunoglobulin (Sigma F1262) diluted 150-fold in KCM, 1% BSA. Slides were washed twice in KCM (5 min. at room temperature), fixed in 4% v/v formaldehyde (10min. at room temperature), rinsed in deionised water and mounted in Vector Shield (Vector lab) supplemented with DAPI (Sigma) at 2μg/ml. Slides were visualized on a Zeiss Axioplan 2 epifluorescence microscope.

### RT-PCR

RNA was extracted from elutriated LCL fractions using the RNeasy mini kit (Qiagen), including on column DNase I digestion, and converted to cDNA using Tetro cDNA synthesis kit (Bioline) according to the manufacturer’s instructions. Realtime RT-PCR was carried out on an Applied Biosystems 7900HT using SensiMix SYBR green master mix (Bioline). 300nM each primer were added to 10μl reactions in 384-well plates. Primers were designed to amplify unspliced pre-mRNA. Primer sequences are given in supplementary table S3. Experiments were carried out in triplicate and 3 technical replicates were set up for each qPCR reaction.

### Gene Expression Microarray

LCL cells were harvested by centrifugation. HeLa cells were shaken off in mitosis as above, replated and grown for a further 6 hours (late G_1_) before harvesting by trypsinisation. RNA was extracted and purified using the RNeasy kit with DNase digestion (Qiagen) according to the manufacturer’s instructions. Double stranded cDNA was synthesised using the cDNA Synthesis System (Roche-Nimblegen), including RNase I and Proteinase K treatment followed by DNA clean up using the PCR purification kit (Qiagen). Samples were labelled with cy3 using a One-Colour Labeling Kit (Roche-Nimblegen), mixed with alignment oligos and sample tracking control oligos (Nimblegen Hybridisation and Sample Tracking Control Kits) and hybridised to a 12×135k HD2 expression array (Roche Nimblegen, containing 3 probes per sequence for 44,049 human sequences) and scanned on a Nimblegen MS200 Microarray scanner. Data were extracted using DEVA (Roche Nimblegen) and normalised by robust multichip average in R. [79]. Ontology analysis was carried out using DAVID [80].

## Availability of data and materials

All expression microarray data and ChIP-seq data is in the process of being submitted to public repositories – Gene expression data in GEO and ChIP-sequencing in SRA - and will be publicly available prior to publication.

## Authors’ contributions

JAH designed and carried out experiments, analysed the data and wrote the manuscript, SA analysed the data, FK analysed the data, CER carried out experiments, GF carried out sequencing, WR advised on study design and supported sequencing, BMT conceived the study, designed experiments, analysed the data and wrote the manuscript.

## Acknowledgements

We are grateful to Rebecca Sie for technical support. We thank Amanda Fisher for encouraging us to try the centrifugal elutriation approach and Roger Grand for unearthing a suitable rotor.

## SUPPLEMENTARY INFORMATION

**Supplementary Table S1:** Functional enrichment of genes with cell cycle-specific enrichment of H3K9ac at their TSS in LCL cells

**Supplementary Table S2:** Functional enrichment of genes with cell cycle-specific enrichment of H3K9ac at their TSS in HeLa cells

**Supplementary Table S3:** Primers for detecting pre-mRNA expression

**Supplementary Figure S1:** Fractionation of cells by cell cycle phase

**Supplementary Figure S2:** Comparison of G_1_ ChIP-seq with ENCODE

**Supplementary Figure S3:** Cell-cycle specific H3K9ac karyotypes in LCL and HeLa cells

**Supplementary Figure S4:** Cell-cycle specific H3K4me3 karyotype in LCL cells

**Supplementary Figure S5:** Cell-cycle specific H3K27me3 karyotype in LCL cells

**Supplementary Figure S6:** H3K4me3 immunofluorescent staining in Human/Mouse Hybrid containing human chromosome 11

**Supplementary Figure S7:** The effect of reducing read count on correlation by Pearson R

**Supplementary Figure S8:** Correlation of significantly enriched probes in H3K4me3 and H3K9ac in LCL

