## Supplementary Material for "Histone modifications form a cell-type-specific chromosomal bar code that modulates and maintains patterns of gene expression through the cell cycle"

**SUPPLEMENTARY INFORMATION**

**Supplementary Table S1:** Functional enrichment of genes with cell cycle-specific enrichment of H3K9ac at their TSS in LCL cells

**Supplementary Table S2:** Functional enrichment of genes with cell cycle-specific enrichment of H3K9ac at their TSS in HeLa cells

**Supplementary Table S3:** Primers for detecting pre-mRNA expression

**Supplementary Figure S1:** Fractionation of cells by cell cycle phase

**Supplementary Figure S2:** Comparison of $G_1$ ChIP-seq with ENCODE

**Supplementary Figure S3:** Cell-cycle specific H3K9ac karyotypes in LCL and HeLa cells

**Supplementary Figure S4:** Cell-cycle specific H3K4me3 karyotype in LCL cells

**Supplementary Figure S5:** Cell-cycle specific H3K27me3 karyotype in LCL cells

**Supplementary Figure S6:** H3K4me3 immunofluorescent staining in Human/Mouse Hybrid containing human chromosome 11

**Supplementary Figure S7:** The effect of reducing read count on correlation by Pearson R

**Supplementary Figure S8:** Correlation of significantly enriched probes in H3K4me3 and H3K9ac in LCL

**Supplementary Table S1: Functional enrichment of genes with cell cycle-specific enrichment of H3K9ac at their TSS or in the gene body in LCL cells.**

Ontology analysis was carried out using DAVID. n: number of genes with significantly enriched H3k9ac at their TSS (-500 to +750, P<0.5) or gene body (+750 to +1750, P<0.05); Count: number of significant genes with specific annotation; %: proportion of genes in significant gene list with specific annotation; P: P-value; FE: Fold enrichment

#### H3K4me3 TSS $G_1$>$G_2$M (n=316)

| Category | Term | Count | % | P | FE |
|---|---|---|---|---|---|
| GOMF | protein tyrosine kinase activity | 21 | 3 | $1.9 \times 10^{-5}$ | 3.0 |
| GOBP | osteoblast differentiation | 8 | 1 | 0.002 | 4.5 |
| GOCC | focal adhesion | 13 | 2 | $9.0 \times 10^{-4}$ | 3.1 |
| GOCC | actin cytoskeleton | 22 | 3 | 0.003 | 2.0 |
| GOBP | regulation of organelle organization | 24 | 3 | $4.1 \times 10^{-5}$ | 2.6 |
| GOBP | regulation of apoptosis | 50 | 7 | 0.005 | 1.5 |

#### H3K4me3 TSS $G_2$M>$G_1$ (n=216)

| Category | Term | Count | % | P | FE |
|---|---|---|---|---|---|
| GOBP | M phase | 22 | 11 | $4.0 \times 10^{-11}$ | 6.2 |
| GOCC | nuclear lumen | 32 | 17 | $1.5 \times 10^{-5}$ | 2.3 |
| GOBP | regulation of cell cycle | 13 | 7 | $2.2 \times 10^{-4}$ | 3.7 |
| GOCC | nucleoplasm | 20 | 10 | $8.2 \times 10^{-4}$ | 2.3 |
| GOBP | spindle organization | 5 | 3 | 0.001 | 10.4 |
| GOCC | condensed chromosome | 7 | 4 | 0.002 | 5.5 |
| GOBP | M phase of meiotic cell cycle | 6 | 3 | 0.004 | 5.7 |

#### H3K9ac TSS $G_1$>$G_2$M (n=124)

| Category | Term | Count | % | P | FE |
|---|---|---|---|---|---|
| GOMF | purine nucleotide binding | 20 | 17 | 0.01 | 1.8 |
| GOMF | ATPase activity | 6 | 5 | 0.04 | 3.1 |

#### H3K9ac TSS $G_2$M>$G_1$ (n=161)

| Category | Term | Count | % | P | FE |
|---|---|---|---|---|---|
| GOCC | nuclear lumen | 23 | 17 | $3.0 \times 10^{-4}$ | 2.3 |
| GOCC | microtubule cytoskeleton | 14 | 10 | $1.0 \times 10^{-4}$ | 3.6 |
| GOCC | spindle | 9 | 7 | $8.1 \times 10^{-6}$ | 8.7 |
| GOBP | mitotic cell cycle | 10 | 7 | 0.002 | 3.4 |
| GOBP | M phase | 11 | 8 | $2.5 \times 10^{-4}$ | 4.2 |

#### H3K9ac Body $G_2$M>$G_1$ (n=26)

| Category | Term | Count | % | P | FE |
|---|---|---|---|---|---|
| GOBP | mitotic cell cycle | 10 | 37 | $3.0 \times 10^{-10}$ | 19.2 |
| GOBP | M phase | 9 | 33 | $4.0 \times 10^{-9}$ | 19.5 |
| GOBP | cell division | 6 | 22 | $3.2 \times 10^{-5}$ | 14.5 |
| GOCC | spindle | 6 | 22 | $1.9 \times 10^{-6}$ | 26.1 |

**Supplementary Table S2: Functional enrichment of genes with cell cycle-specific enrichment of H3K9ac at their TSS in HeLa cells.**

Ontology analysis was carried out using DAVID. n: number of genes with significantly enriched TSS H3K9ac (P<0.5); Count: number of significant genes with specific annotation; %: proportion of genes in significant gene list with specific annotation; P: P-value; FE: Fold enrichment

**$G_1$>$G_2$M (n=547)**

| Category | Term | Count | % | P | FE |
|---|---|---|---|---|---|
| GOMF | nucleoside binding | 86 | 17 | $7.0x10^{-11}$ | 2.0 |
| GOBP | cell cycle phase | 27 | 5 | $5.5x10^{-5}$ | 2.4 |
| GOCC | chromosome | 25 | 5 | 0.002 | 2.0 |
| GOMF | cofactor binding | 18 | 4 | $2.8x10^{-4}$ | 2.8 |

**$G_2$M>$G_1$ (n=315)**

| Category | Term | Count | % | P | FE |
|---|---|---|---|---|---|
| GOCC | cytosolic ribosome | 23 | 8 | $1.1x10^{-21}$ | 18.0 |
| GOCC | nuclear lumen | 48 | 17 | $8.8x10^{-7}$ | 2.1 |
| GOBP | mitosis | 11 | 4 | 0.004 | 3.0 |

**$G_1$>M (n=1552)**

| Category | Term | Count | % | P | FE |
|---|---|---|---|---|---|
| GOMF | cytoskeletal protein binding | 66 | 5 | $1.0x10^{-5}$ | 1.7 |
| GOBP | protein processing | 20 | 1 | $6.1x10^{-4}$ | 2.4 |
| GOMF | nucleoside binding | 163 | 12 | $3.6x10^{-5}$ | 1.3 |
| GOMF | GTPase regulator activity | 55 | 4 | $2.1x10^{-5}$ | 1.8 |
| GOMF | amine binding | 19 | 1 | 0.001 | 2.3 |
| GOMF | SH3/SH2 adaptor activity | 11 | 1 | 0.003 | 3.0 |
| GOBP | cytoskeleton organization | 49 | 3 | 0.004 | 1.5 |

**M>$G_1$ (n=1556)**

| Category | Term | Count | % | P | FE |
|---|---|---|---|---|---|
| GOBP | translational elongation | 75 | 5 | $8.9x10^{-59}$ | 9.1 |
| GOCC | intracellular non-membrane-bounded organelle | 297 | 21 | $7.1x10^{-14}$ | 1.5 |
| GOBP | ribosome biogenesis | 33 | 2 | $1.9x10^{-9}$ | 3.3 |
| GOCC | nuclear lumen | 159 | 11 | $3.6x10^{-6}$ | 1.4 |
| GOBP | mRNA processing | 39 | 3 | 0.01 | 1.5 |
| GOBP | biological adhesion | 83 | 6 | $5.2x10^{-4}$ | 1.4 |
| GOCC | heterogeneous nuclear ribonucleoprotein complex | 11 | 1 | $8.9x10^{-8}$ | 8.3 |
| GOBP | response to metal ion | 21 | 2 | 0.004 | 2.0 |
| GOBP | regulation of cellular protein metabolic process | 60 | 4 | $7.0x10^{-4}$ | 1.5 |
| GOCC | membrane-bounded vesicle | 64 | 5 | 0.002 | 1.5 |
| GOBP | cell cycle | 94 | 7 | $9.3x10^{-5}$ | 1.5 |
| GOBP | mitotic cell cycle | 47 | 3 | 0.003 | 1.6 |

**$G_2M$>M (n=822)**

| Category | Term | Count | % | P | FE |
|---|---|---|---|---|---|
| GOBP | regulation of transcription from RNA polymerase II promoter | 59 | 8 | $2.2x10^{-7}$ | 2.1 |
| GOBP | cell motion | 37 | 5 | $1.4x10^{-4}$ | 2.0 |
| GOBP | response to hormone stimulus | 29 | 4 | $6.8x10^{-4}$ | 2.0 |
| GOBP | negative regulation of gene expression | 39 | 5 | $1.0x10^{-4}$ | 2.0 |
| GOBP | regionalization | 21 | 3 | $1.0x10^{-4}$ | 2.7 |
| GOBP | tube development | 24 | 3 | $2.0x10^{-5}$ | 2.8 |
| GOCC | cell junction | 39 | 5 | $5.9x10^{-5}$ | 2.0 |
| GOBP | embryonic morphogenesis | 26 | 3 | $5.0x10^{-4}$ | 2.1 |
| GOMF | metal ion binding | 202 | 27 | $5.1x10^{-4}$ | 1.2 |
| GOBP | Wnt receptor signaling pathway | 15 | 2 | $7.7x10^{-4}$ | 2.9 |
| GOBP | cell motion | 37 | 5 | $1.4x10^{-4}$ | 2.0 |
| GOMF | GTPase regulator activity | 33 | 4 | $1.8x10^{-4}$ | 2.0 |
| GOBP | actin cytoskeleton organization | 19 | 3 | 0.004 | 2.1 |
| GOBP | cell adhesion | 46 | 6 | $8.8x10^{-4}$ | 1.7 |
| GOBP | regulation of phosphorylation | 39 | 5 | $1.8x10^{-5}$ | 2.1 |
| GOMF | transcription repressor activity | 24 | 3 | 0.004 | 1.9 |

**M>$G_2M$ (n=1490)**

| Category | Term | Count | % | P | FE |
|---|---|---|---|---|---|
| GOBP | translational elongation | 62 | 5 | $1.8x10^{-42}$ | 8.0 |
| GOCC | small ribosomal subunit | 28 | 2 | $2.2x10^{-14}$ | 5.7 |
| GOCC | large ribosomal subunit | 28 | 2 | $1.4x10^{-13}$ | 5.4 |
| GOCC | intracellular non-membrane-bounded organelle | 301 | 22 | $3.9x10^{-15}$ | 1.5 |
| GOMF | cytoskeletal protein binding | 64 | 5 | $8.1x10^{-5}$ | 1.6 |
| GOCC | cytoskeleton | 151 | 11 | $6.2x10^{-6}$ | 1.4 |
| GOCC | nuclear lumen | 149 | 11 | $1.8x10^{-4}$ | 1.3 |
| GOBP | RNA processing | 63 | 5 | 0.001 | 1.5 |
| GOBP | cytoskeleton organization | 56 | 4 | $1.6x10^{-4}$ | 1.7 |
| GOBP | cell adhesion | 71 | 5 | 0.01 | 1.3 |
| GOCC | cell cortex | 23 | 2 | 0.002 | 2.0 |

### Supplementary Table S3: Primers for detecting pre-mRNA expression

Primers were designed such that one primer binds in an exon and the other in the adjoining intron so as to detect unspliced pre-mRNA.

| Gene | Forward Primer | Reverse Primer |
|---|---|---|
| **ZBP1** | ggcagagaaggtacagtact | atcatgatctggaccaagcg |
| **SKA3** | gtcatatgtgatctgtgattgacc | ctcattggataatcttcaaagtcttc |
| **PRDM2** | caaactctacagttctgtatgtggt | gaaacttctttgattacatgaaccc |
| **MEIS2** | ccacaccctccaaacttgttg | tcgcacttctcaaagaccaga |
| **NBN** | gtctagcagccccggttac | gacgcgacgcaggaatca |
| **CCNB1** | gacttactcgagccttcgtgg | aagagctgttcttggcctcag |
| **CCNF** | gtggcacagggaatagtaacca | ttggcacacctacagtggacc |
| **CDCA3** | ttgagccacgagctgttgtgc | tgtgactgggacgctcttggc |
| **DLGAP5** | gtttcacgctagaaggtggc | ctcgcattatcagcttcggg |
| **HIF20A** | gcgactaggtgtgagtaagcc | gcagctccgatttcttaaagcc |
| **PLK1** | acaacgacttcgtgttcgtgg | ttgctgggtactctttccagc |
| **TTK** | tttccccagcgcagctttctg | tttagccgcgtcgcatacctg |

**Supplementary Figure S1: Fractionation of cells by cell cycle phase**

Flow cytometry analysis using propidium iodide staing for DNA content of cells following cell cycle fractionation by FACS (A), centrifugal elutriation (B) and mitotic shake off (C) of HeLa cells in culture (D), showing mostly adherent cells and rounded, loosely attached mitotic cells. (E) Chromatin from $G_1$ and $G_2M$ cells show similar fragment size and distribution following micrococcal nuclease digestion, DNA extraction and 2% agarose gel electrophoresis (S=supernatant, P=pellet following dialysis of digested chromatin).

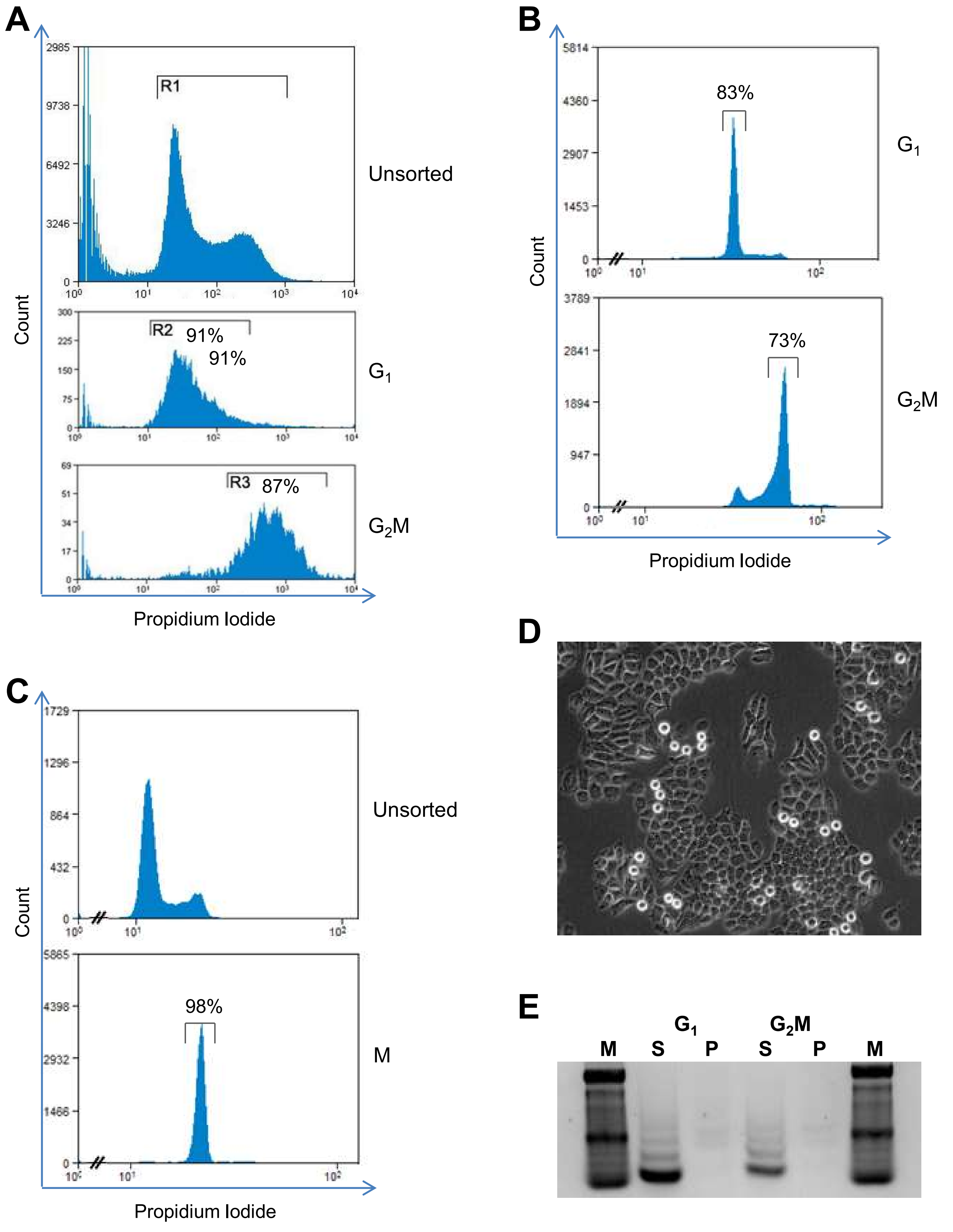

**Supplementary Figure S2: Comparison of $G_1$ ChIP-seq with data from ENCODE**

We compared our ChIP from acetone-fixed, FACS-sorted $G_1$ LCLs with data from ENCODE for the LCL line GM12878 (formaldehyde fixed, asynchronous). Genome wide comparisons were carried out using 5000bp rolling window probes. A) Scatter plots for H3K9ac, H3K4me3 and H3K27me3 are shown. Grey dots represent all 5000bp probes, blue dots represent specific features as shown. Strong positive correlations were observed for all three modifications. TSS: transcription start site, CpG: CpG island. B) Pearson R values for correlations of probes overlapping specific genomic features, red highlight: R > all probes, green: R < all probes. For H3K9ac and H3K4me3, the strength of correlation increased at gene regulatory regions, transcription start sites (R=0.905 and 0.854 respectively) and CpG islands. C) Screenshot of the HoxA cluster displayed in SeqMonk comparing ChIP-seq signals from our $G_1$ sorted LCLs and the ENCODE LCL GM12878.

**A**

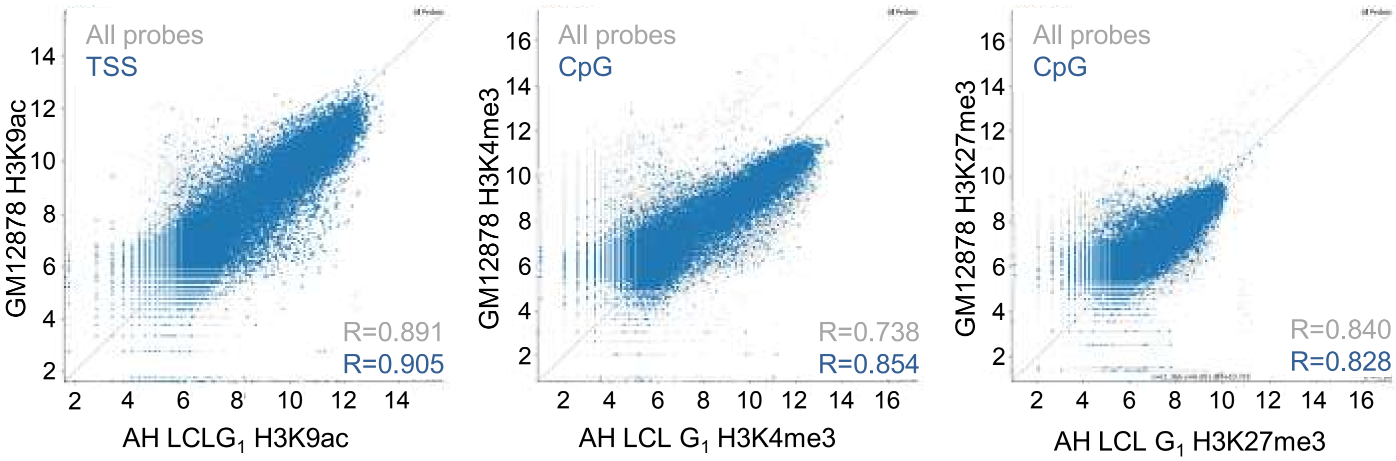


**B**

| Pearson R | H3K9ac | H3K4me3 | H3K27me3 |
|---|---|---|---|
| **All probes** | 0.891 | 0.738 | 0.840 |
| **TSS** | 0.905 | 0.854 | 0.770 |
| **CpG Islands** | 0.924 | 0.906 | 0.828 |
| **Genes** | 0.806 | 0.707 | 0.724 |

**C**

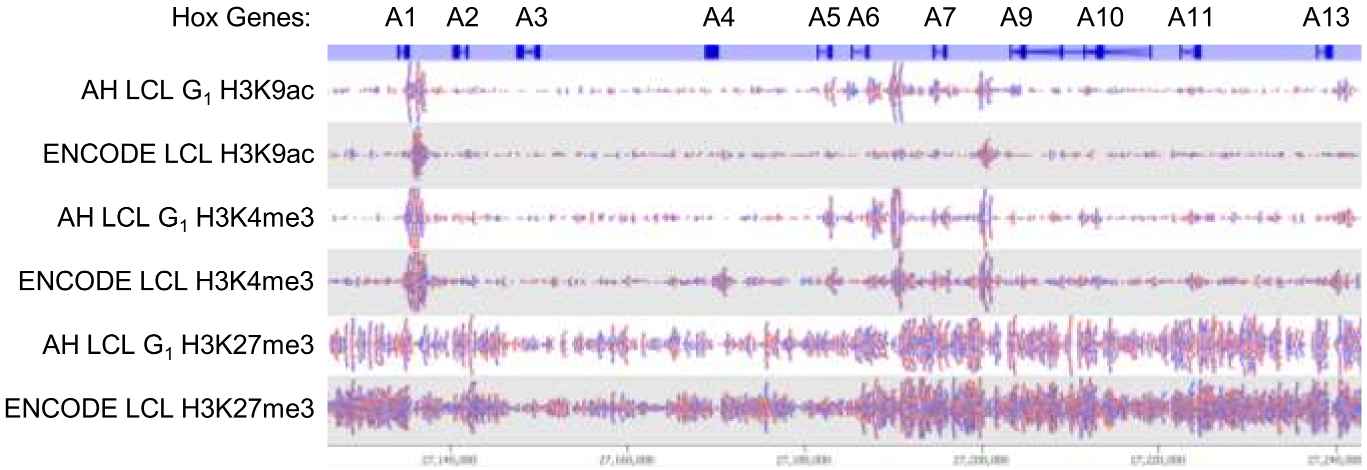

**Supplementary Figure S3: Cell-cycle specific H3K9ac karyotypes in LCL and HeLa cells.** ChIPseq data was analysed using 700bp rolling windows and displayed as blocks using a red-green colour scale in SeqMonk. Maximum reolution (1 pixel) is approximately 1.5MB. $G_1$ and $G_2M$ fractions were sorted by FACs (LCL, **A**) or centrifugal elutriation (HeLa, **B**). LCL ChIP-seq karyotypes are shown aligned with metaphase chromosomes stained with antibody to H3K9ac as described in Terrenoire et al. (2010). Mitotic HeLa cells were harvested by mitotic shake-off.

### A: H3K9ac in LCL

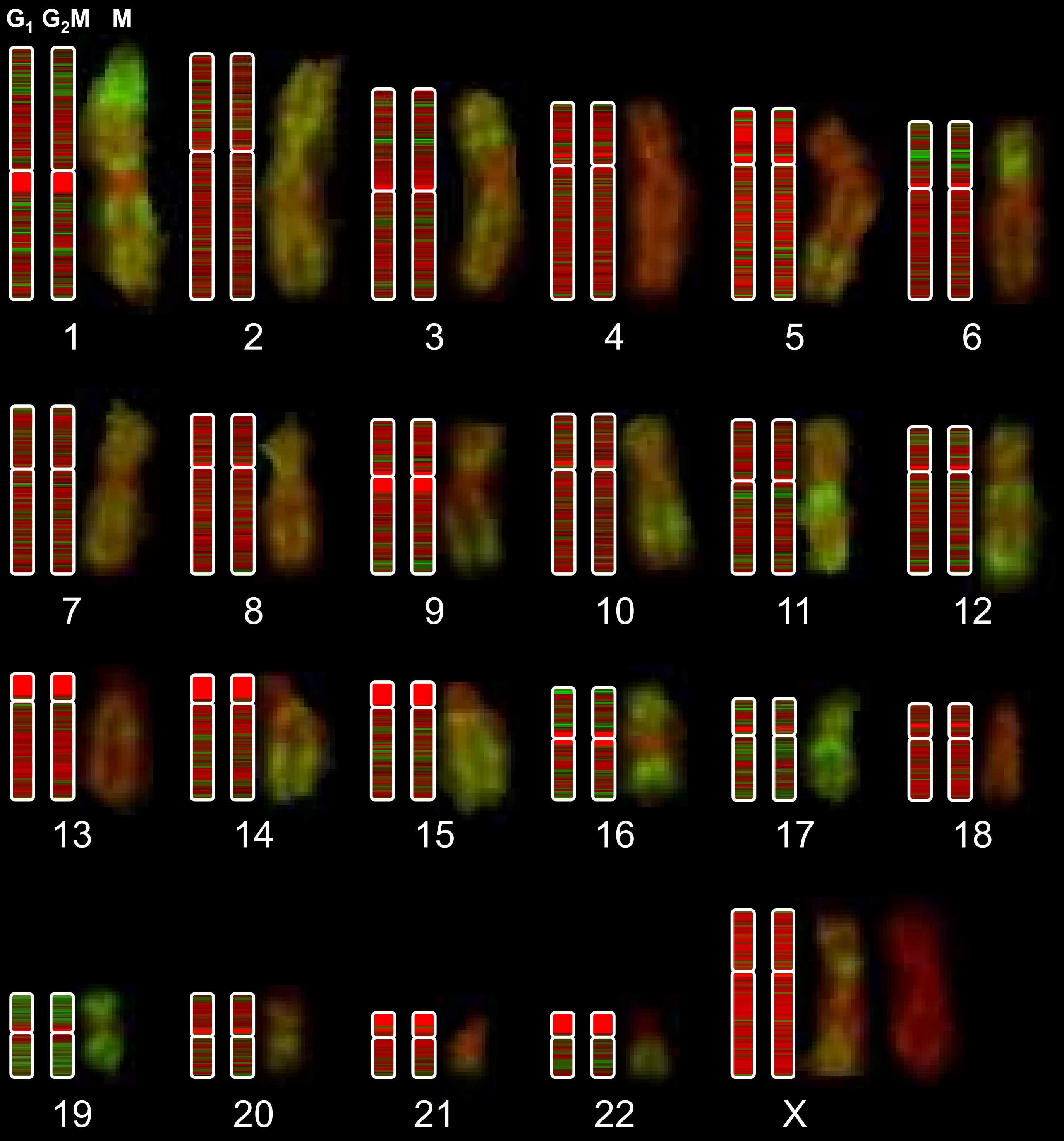

**B: H3K9ac in HeLa**

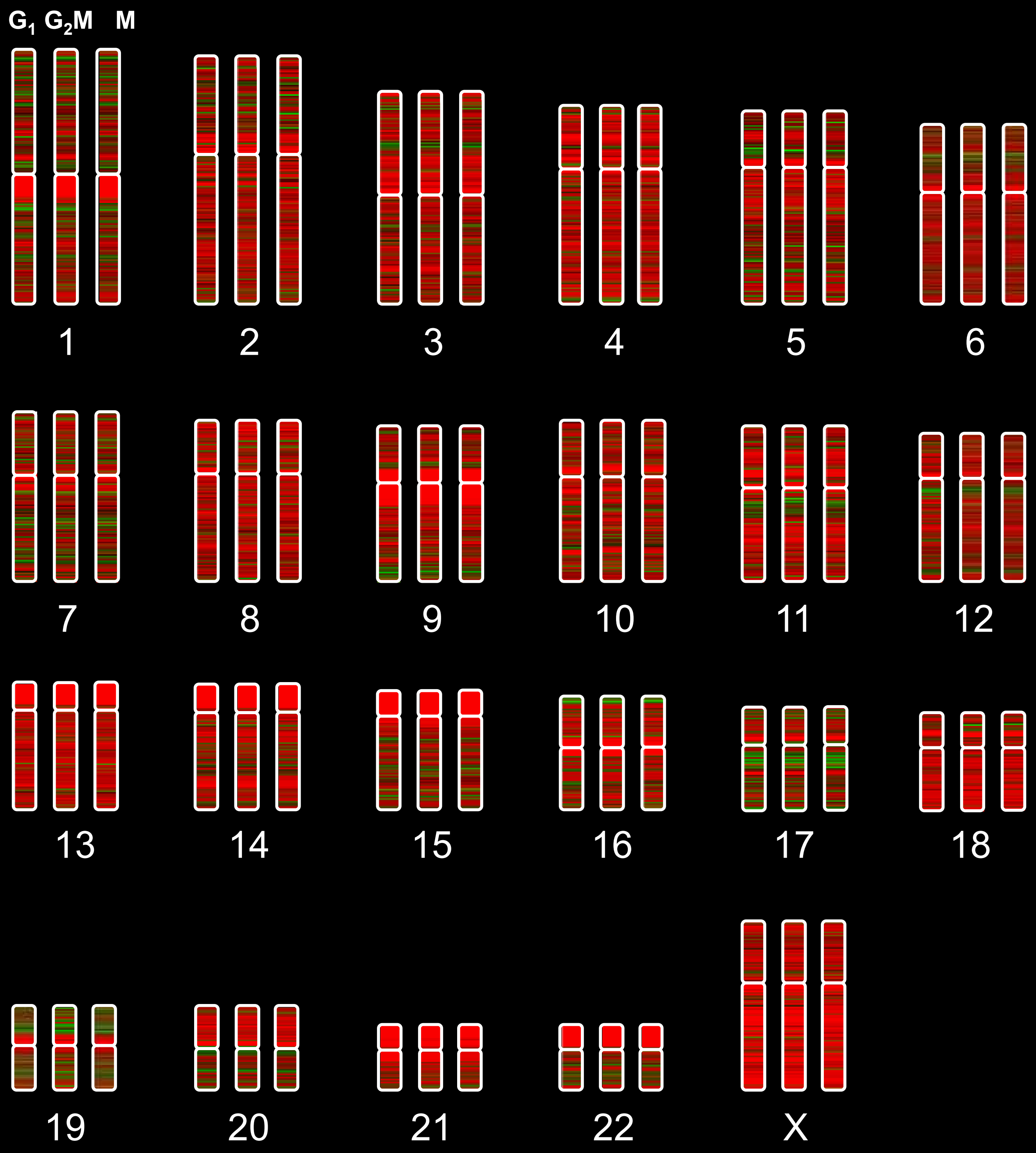

**Supplementary Figure S4: Cell-cycle specific H3K4me3 karyotype in LCL cells.** ChIPseq data was analysed using 700bp rolling windows and displayed as bars using a red-green colour scale in SeqMonk. Maximum reolution (1 pixel) is approximately 1.5MB. $G_1$ and $G_2M$ fractions were sorted by FACs. ChIP-seq karyotypes are shown aligned with metaphase chromosomes stained with antibody to H3K4me3 as described in Terrenoire et al. (2010).

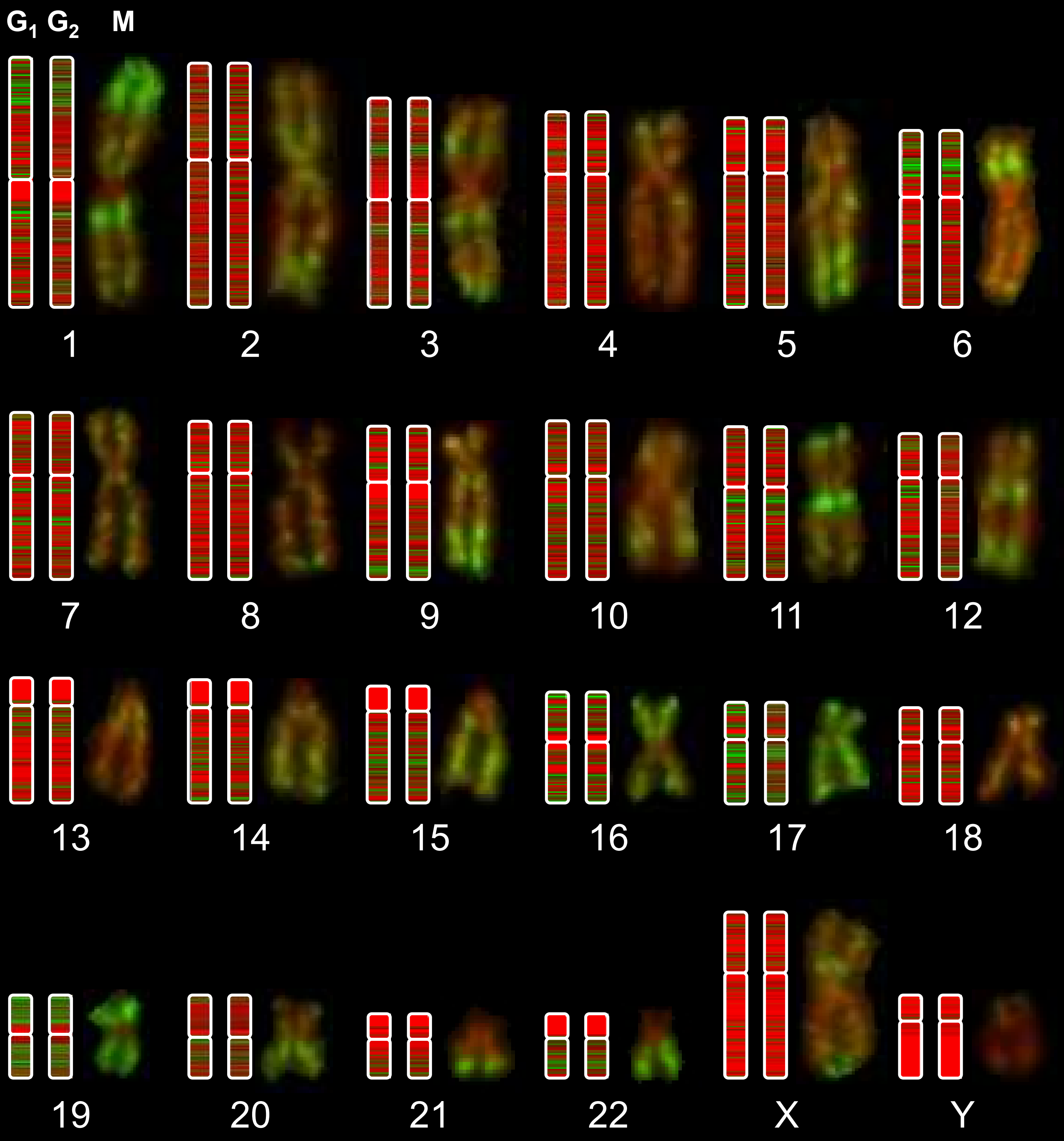

**Supplementary Figure S5:** Cell-cycle specific H3K27me3 karyotype in LCL cells. ChIPseq data was analysed using 700bp rolling windows and displayed as bars using a red-green colour scale in SeqMonk. Maximum reolution (1 pixel) is approximately 1.5MB. $G_1$ and $G_2M$ fractions were sorted by FACs. ChIP-seq karyotypes are shown aligned with metaphase chromosomes stained with antibody to H3K27me3 as described in Terrenoire et al. (2010).

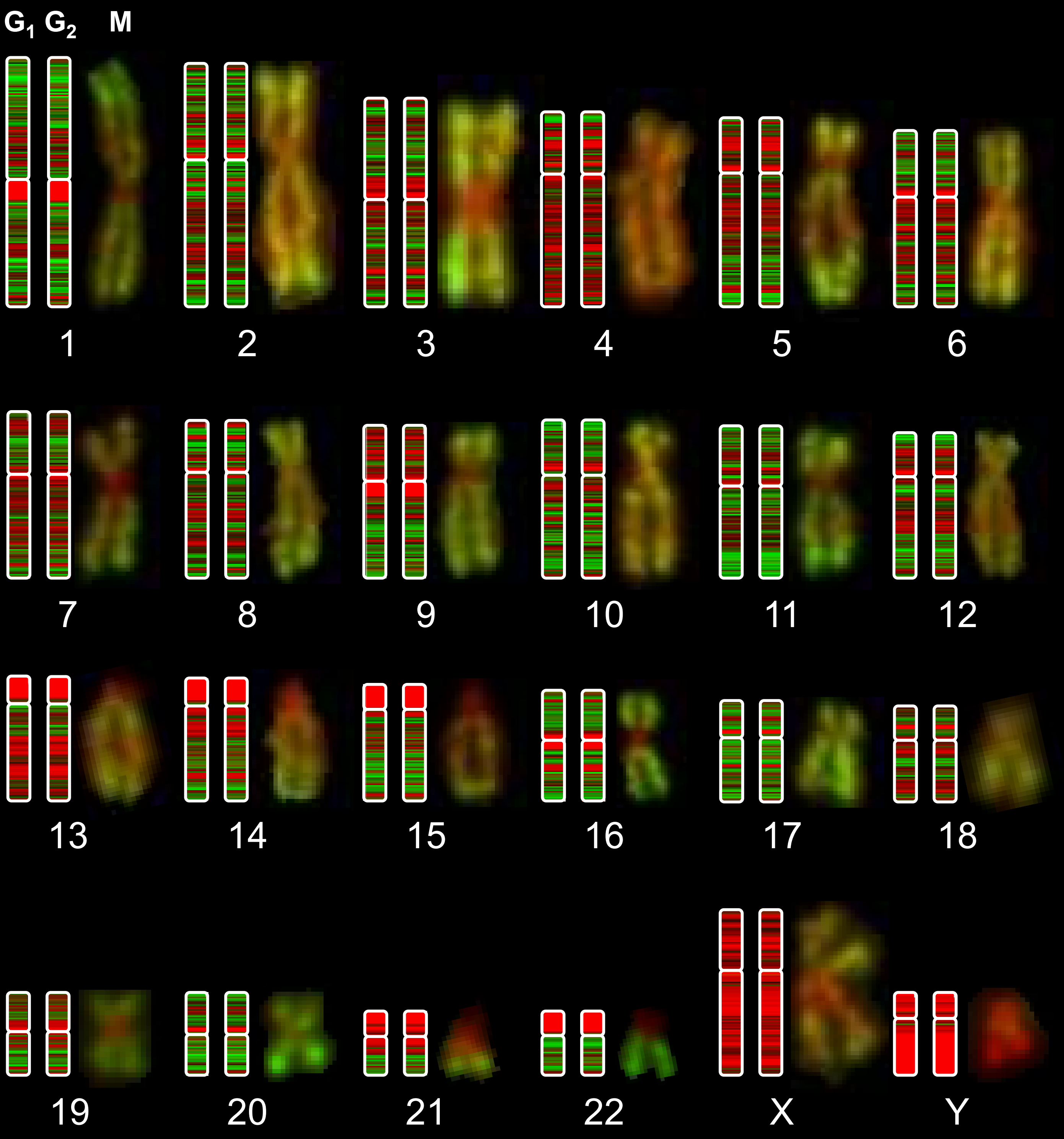

**Supplementary Figure S6:** H3K4me3 immunofluorescent staining in Human/Mouse Hybrid containing human chromosome 11

**A:** Metaphase spread from the mouse/human hybrid line GM11941, created by fusing human lymphocytes to mouse 1R cells and containing a complete human chromosome 11, stained with antibody to H3K4me3 (green) and dapi (red). Human chromosome 11 is easily identifiable as submetacentric in comparison to the telocentric mice chromosomes. Human chromosome 11 is marked with an arrow and magnified (inset).

**B:** H3K4me3 ChIP-seq signal and immunofluorescent staining of chromosome 11 in LCL (from Supplementary figure S4), included for comparison.

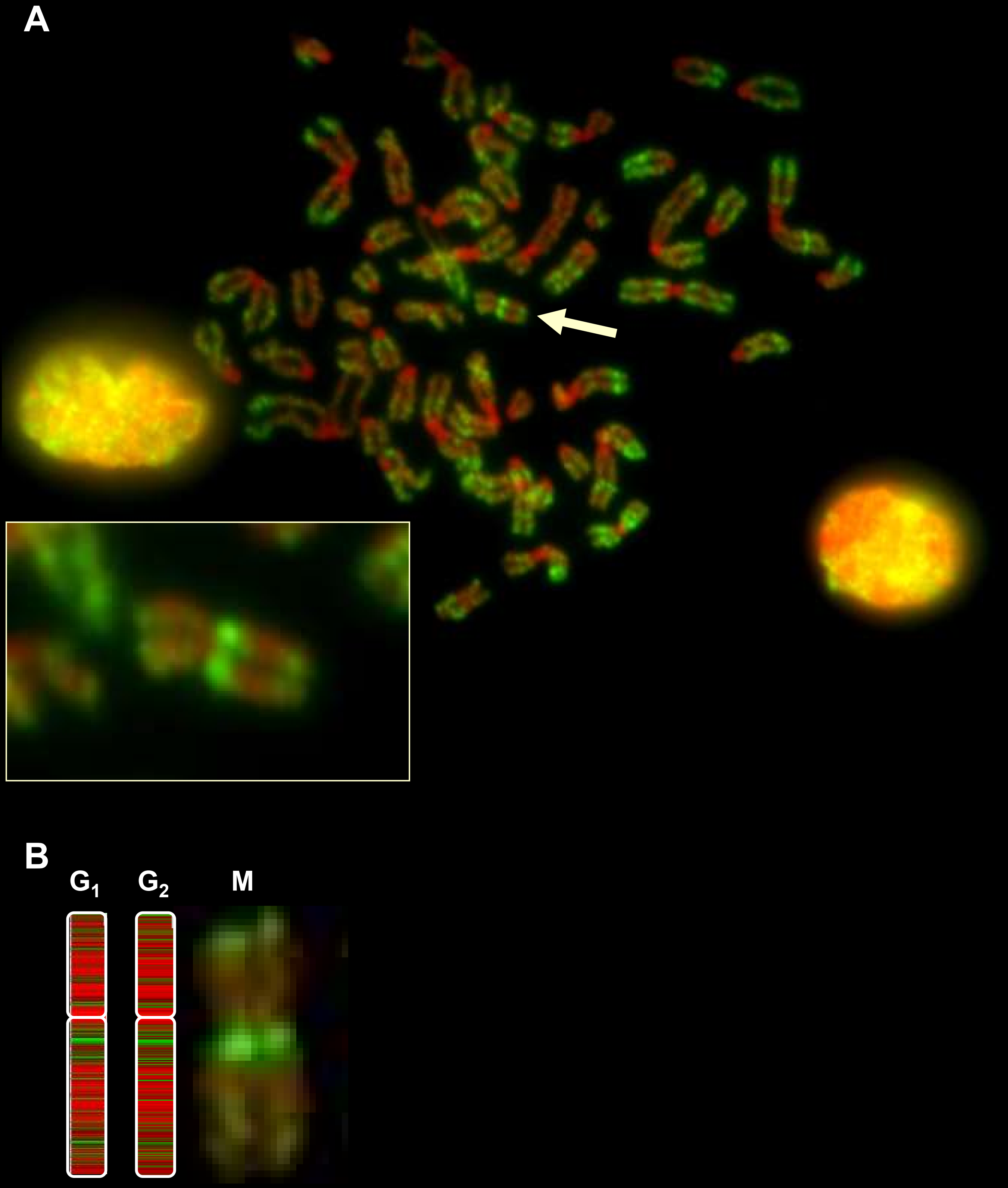

**Supplementary Figure S7:** The effect of reducing read count on correlation by Pearson R

A ChIP-seq data set of H3K9ac in G1 in LCL (17.5M reads) was sampled to produce datasets with reduced read counts as indicated. These down-sampled datasets were then compared to the original using rolling windows of various sizes. The comparison of the complete datasets for H3K9ac in $G_1$ vs $G_2M$ is shown for reference.

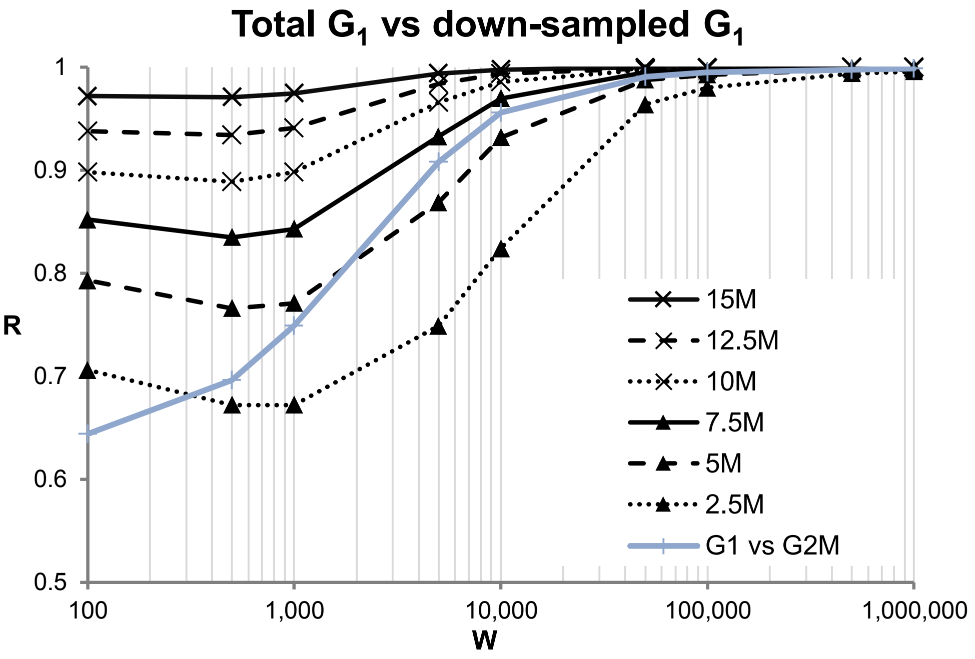

**Supplementary Figure S8:** Correlation of significantly enriched probes in H3K4me3 and H3K9ac in LCL. **A: :** Scatterplot showing correlation of H3K4me3 at TSS (as in figure 4B) with probes significantly enriched in H3K9ac in $G_1$ (blue) or $G_2M$ (red) highlighted. **B:** Scatterplot showing correlation of H3K9ac at TSS (as in figure 4D) with probes significantly enriched in H3K4me3 in $G_1$ (blue) or $G_2M$ (red) highlighted.

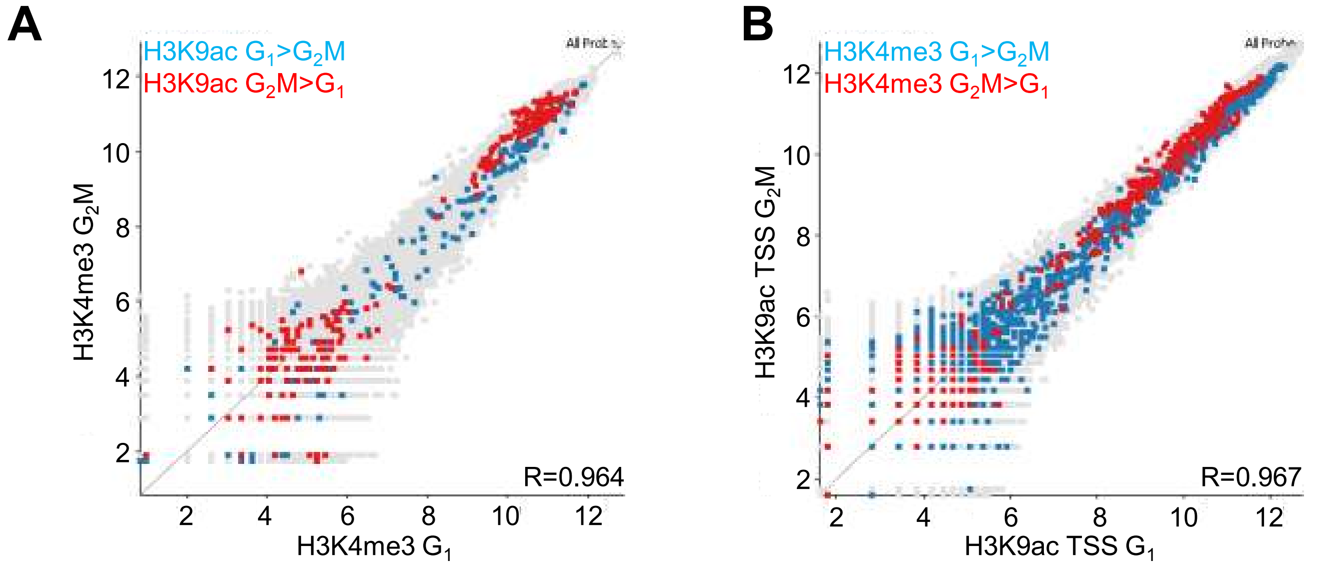
